# Localised Astrocyte Ca2+ Activity Regulates Neurovascular Coupling Responses to Active Sensing

**DOI:** 10.1101/2024.04.16.589720

**Authors:** Jakob Akbar Stelzner, Aske Krogsgaard, Gabriele Kulkoviene, Leonora Sperling, Barbara Lykke Lind

## Abstract

Neurovascular coupling (NVC) ensures sufficient and targeted blood flow during increased neuronal activity. Astrocytic participation in NVC has long been debated, likely due to the intricacy of the intracellular Ca2+ fluxes and the diversity of their regulatory capacities. As astrocyte signaling changes with brain states, we focused on their involvement in voluntary sensing in freely behaving mice. We used 2-photon microscopy to record cellular and vascular activity in the whisker barrel cortex of awake head-fixed animals. The NVC initiated by volitional whisking in the resting mouse was compared to the whisking preceding locomotion and experimenter-evoked whisker deflections. We developed an analysis method to detect early, subcellular astrocytic activity and found it corresponded with neuronal and vascular responses under all three conditions. After the depletion of noradrenaline (NA), the early astrocytic Ca2+ response to volitional whisking was only moderately reduced and primarily in astrocytic processes closest to the blood vessels. Meanwhile, the dilation of 1^st^ order capillaries was also reduced. Together, these findings demonstrate significant disruptions in the focal regulation of cerebral blood flow, potentially limiting the sustenance of activated neurons. This disruption appeared to translate into behavioral aberrations, as NA-depleted mice exhibited an extended period of exploratory whisking prior to locomotion. Remarkably, NA-depletion did not alter cellular or blood flow responses to locomotion or experimenter-evoked whisking. Our study confirms an astrocytic contribution to NVC, which is relevant during volitional sensing. It also suggests that self-directed sensory processing depends on an appropriate NVC response, which itself depends on NA and astrocyte activity.

## Introduction

Neurovascular coupling (NVC) is the process that recruits additional local cerebral blood flow (CBF) to match the increased energy demands during local brain activation (Roy and Sherrington 1890). Over the years, several cell types have been identified to partake in this process, amongst them astrocytes, which are known to possess various, and even counteracting, regulatory mechanisms (Howarth, Mishra et al. 2021, Schaeffer and Iadecola 2021). It is not established when and where each of the regulatory mechanisms contributes to blood flow recruitment. Disentangling the regulation of NVC has significant clinical relevance, as many CNS diseases present with disruptions to NVC (Schaeffer and Iadecola 2021). We know that the dynamics of cortical excitation in awake and freely behaving animals depend on the shifting activation of complex neuronal circuits (Petersen 2019) and on neuromodulator release (Collins, Francis et al. 2023). These different activity patterns depend on and contribute to both global shifting brain states and localized responses, with may give a high regional diversity in NVC responses (Drew 2022). We propose that this diversity depends on the context of neuronal excitation. If this is the case, then the behavioral state influences which blood flow regulating mechanisms are at work as neurons are excited. The implication would be that additional energy substrates and oxygen are not provided to a similar degree or through identical mechanisms during different cognitive tasks.

Astrocytes are ideally situated for integrating information about synaptic activity and relaying this information to blood vessels as their processes enwrap both synapses and the brain microvasculature (Mathiisen, Lehre et al. 2010, McCaslin, Chen et al. 2011). However, investigations into the astrocytic contribution to NVC have reached different conclusions (Takano, Tian et al. 2006, Petzold, Albeanu et al. 2008, Nizar, Uhlirova et al. 2013, Bonder and McCarthy 2014, Institoris, Rosenegger et al. 2015, Lacroix, Toussay et al. 2015, Masamoto, Unekawa et al. 2015, Del Franco, Chiang et al. 2022). We and others have previously shown that astrocytic Ca2+ transients may occur at fast timescales, comparable to neuronal activities and that these fast transients may be involved in the dilatory phase of NVC(Lind, Brazhe et al. 2013, Otsu, Couchman et al. 2015, Lind, Jessen et al. 2018). Other roles for astrocytes have been suggested, such as response amplification (Gu, Chen et al. 2018, Institoris, Vandal et al. 2022) or imposing the late vasoconstrictive phase (Mulligan and MacVicar 2004, Metea and Newman 2006, Gordon, Choi et al. 2008, Girouard, Bonev et al. 2010). Astrocytic activity depends on the brain state (Bindocci, Savtchouk et al. 2017, Bojarskaite, Bjornstad et al. 2020, Krogsgaard, Sperling et al. 2023, Reitman, Tse et al. 2023). Hence, the ambiguous observations may be due to different behavioral contexts in the studies, the most prominent being anesthesia. Another difference is in where and how astrocytic activity is identified and analyzed. We now know that much astrocytic activity is contained within sub-cellular structures(Bindocci, Savtchouk et al. 2017). In addition, the astrocytic contribution to NVC may be capillary-specific(Biesecker, Srienc et al. 2016, Mishra, Reynolds et al. 2016). Hence, we focused our investigation on the cells surrounding the vascular inflow tract where a 1^st^ order capillary branches from the penetrating arteriole (PA). PA expansions provide an increased inflow to deeper cortical layers, whereas capillary dilation increases local blood flow. In this way, a 1^st^ order branch point represents a key regulation point within the vascular tree for the delivery of freshly oxygenated blood to the cerebral parenchyma (Grubb, Cai et al. 2020, Hartmann, Coelho-Santos et al. 2022). In the whisker barrel cortex, these regulated dilations are expected to match the area of neuronal excitation created by sensory input through whisker deflections. We investigated the neuronal and astrocytic Ca2+ activity in the cells surrounding these junctions in the somatosensory barrel cortex during volitional whisking of awake mice.

Whisker deflections activate the whisker barrel cortex through thalamocortical input (Petersen 2019). Up to now, most studies of NVC in the barrel cortex have used a forced stimulation protocol (Ding, O’Donnell et al. 2013, Stobart, Ferrari et al. 2018, Tran, Peringod et al. 2018, Sharma, Gordon et al. 2020). However, this approach only targets one pattern of murine cognition, namely the unsuspected excitation of the sensory cortex. Volition whisker movements are also timed with the activation of noradrenergic projections from the locus coeruleus (LC) (Kim, Jung et al. 2016, Collins, Francis et al. 2023). It has been shown that the release of noradrenaline (NA) from the LC alone can trigger a cortical blood flow response (Toussay, Basu et al. 2013, Lecrux, Sandoe et al. 2017). The degree of NA contribution to NVC could thus depend on whether the whisking is due to a change in the alertness of the animal and part of voluntary spatial surveillance and sensory processing (Shimaoka, Harris et al. 2018). To disentangle the role of these modulatory projections, we divided NVC responses during whisker deflection into three categories, of which two were voluntary: volitional whisking at rest, volitional whisking related to locomotion, and an air puff evoked stimulation. As we speculated that astrocytic contributions to whisking-evoked NVC were context and behavior-dependent, we investigated whether astrocytic activity differs between rest versus locomotion and whether this difference could be partially dependent on cortical NA dynamics.

We found that regardless of the context of whisker deflection, a NVC response in the barrel cortex always entailed a neuronal excitation, shortly followed by an astrocytic Ca2+ response and then a dilation of the neighboring blood vessels. After dilation had peaked, we observed a second astrocytic response. The size and location of the early astrocytic Ca2+ activity correlated with the size and location of neuronal excitation across all behavioral categories. We used DSP-4, a neurotoxin that eliminates the noradrenergic projections from locus coeruleus (Slezak, Kandler et al. 2019, Renden, Institoris et al. 2024) to investigate the NA contribution to NVC. We found that NVC was differentially affected in each of our three whisking-related behavioral categories. In air-puff-evoked NVC, it did not affect the astrocytes or the NVC response. In volitional whisking at rest, the NA depletion affected both astrocytic Ca2+ responses and 1^st^ order capillary dilations. This result indicates a critical contribution from astrocytes to the laminar and localized blood flow regulation within the volitional and anticipated sensory response to whisking. Remarkably, the reduction in whisking-associated astrocyte activity and local blood flow coincided with a delay between the onset of volitional whisking and subsequent locomotion. Once locomotion was initiated, neuronal, blood vessel and early astrocytic activities were no longer inhibited by the lack of NA release from the LC. Hence, our findings show that by using a context-dependent analysis method, it is possible to identify astrocytic activity that is both temporally and spatially recurrent with neuronal activity and necessary for targeted NVC responses to anticipated sensory input.

## Results

### Self-initiated whisking events trigger NVC responses, which depend on whisking duration

We used awake, head-fixed mice to study intracellular Ca2+ activity and hemodynamic responses during natural spontaneous whisking (Figure 1A). Data was recorded with two-photon microscopy imaging (2PM) from layer II/III of the whisker barrel cortex of adult mice. Blood flow regulation was evaluated by staining the lumen of the vasculature with Texas Red dextran, which was given I.V. prior to the imaging. The lumen staining exposed dilations and constrictions of the blood vessels. Viral vectors were injected to express Ca^2+^ indicators to specific cell types. During imaging, the head was fixed beneath the objective, but the mouse was free to move within an airlifted mobile homecage, which tracked the animals’ movements (Figure 1A). The benefit of using the whisker barrel cortex to study neurovascular coupling (NVC) responses was that we had a visual cue to when sensory input flowed via the thalamus to the cortex. We got this by monitoring the snout of the mouse with an IR camera during the two-photon imaging (Figure 1A) and then employing a motion detection algorithm to record whisking events (Sup. Figure S1). Relating these data with the output from the sensor in the homecage, we could detect whether the mouse was either resting or running as it moved its whiskers. We found that locomotion was always associated with whisking, but the resting mouse also regularly moved its whiskers, with a mode of every 2 seconds (Sup. Figure S2). The whisking periods in the resting animal lasted on average 0.96s±0.046 s (Sup. Figure S2), of which 22% were of a >1 s duration. We then investigated how well these volitional whisking events recruited a NVC response in the receiving whisker barrel cortex. The NVC was evaluated at the first branching point in the vascular inflow tract. Here we quantified the cross area of both the penetrating arterioles and the 1^st^ order capillary branching from it (Grubb, Cai et al. 2020, Hartmann, Coelho-Santos et al. 2022) (Figure 1B). There were frequent changes in the lumen size of these vessels, and they appeared to correlate to the volitional whisking activity of the spontaneously behaving mouse. To quantify whether this was part of volitionally triggered neurovascular coupling, we aligned spontaneously occurring whisking events with the changes in the vessel area (Figure 1C). Sorting whisking events at ½ second intervals revealed that the onset of natural whisking in stationary mice was strongly associated with vasodilation of both the PA and 1^st^ order capillary and that even the shortest whisking event would cause a dilation (Figure 1D and E). As would be expected, longer whisking events evoked larger vasodilations of PA and 1^st^ order capillary (Figure 1F) (Pearson correlation amplitude PA: R = 0.98, p = 0.0006 and AUC R = 0.97, p = 0.0012, N=20 FOV in 16 mice, 1^st^ order cap.: R = 0.94, p = 0.0049 and AUC R = 0.84, p = 0.0038 N= 10 FOV in 7 mice). Our results suggest that all dilations, larger or smaller, reflect the sensory stimulus evoked by the whisker’s movements.

**Figure 1.**
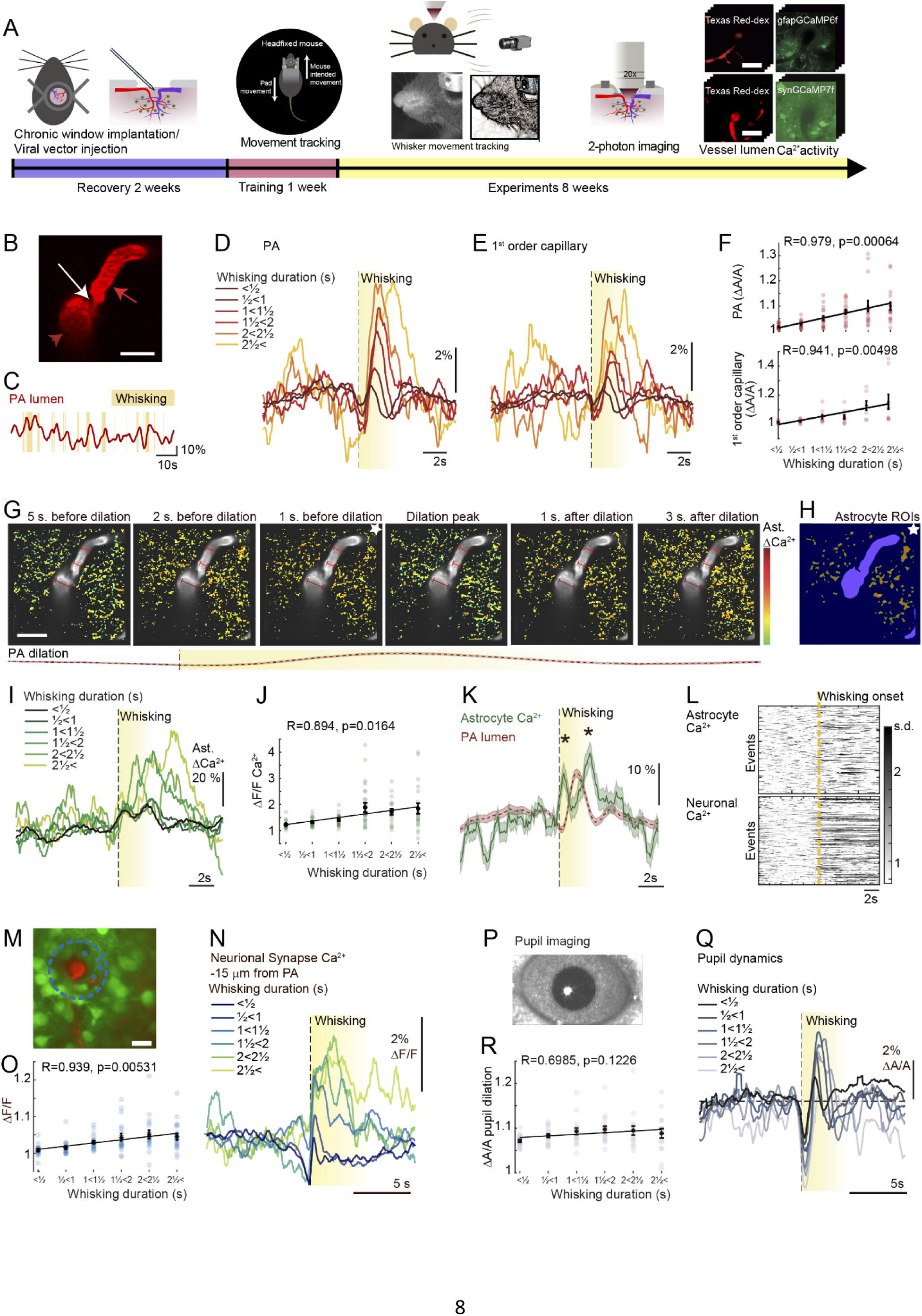
Self-initiated whisking events trigger neurovascular coupling responses that depend on whisking duration and exhibit spatially recurrent rises in neuronal and astrocytic Ca2+ activity before vasodilation. **A)** Experimental setup and protocol. A viral vector was injected when the chronic window was implanted. After two weeks of recovery, mice were gradually habituated to experimental conditions. Locomotion events were measured as the movements of the mobile homecage. Whisker movement was monitored using a video camera and analyzed for natural whisking activity. Cellular Ca2+ activity and vascular responses were measured using two-photon microscopy. Scalebars 40 µm. **B)** Texas Red-dextran loaded penetrating arteriole (PA, *red arrow)* and 1^st^ order capillary (*light red arrow with stalk*) branching point in the layer II/III of whisker barrel cortex. A clear precapillary sphincter is visible (*white arrow*). Scalebar 20µm. **C)** Example traces of PA lumen changes (*red*) overlaid with *yellow* shaded areas representing whisking. **D)** The average normalized lumen of PA is aligned with the onset of all volitional whisking events of increasing duration. N= 20 PA in 16 mice. **E)** The average normalized lumen of 1^st^ order capillary aligned to onset of all volitional whisking events of increasing duration. N= 10 capillaries in 7 mice. **F)** <u>Top</u>: Average max amplitude of PA vasodilations as a function of whisking duration. Pearson correlation R = 0.98, p = 0.0006. N= 20 PA in 16 mice. <u>Bottom</u>: Average max amplitude of 1^st^ order capillary vasodilations as a function of whisking duration. Pearson correlation R = 0.94, p = 0.004. N= 10 capillaries in 7 mice. **G)** Averaged and normalized 2-photon images from a FOV 120µm below the cortical surface, aligned to dilation peak associated with volitional whisking (Outlined below). Averaged over 1 s intervals. Ca2+ activity normalized to s.d. values per pixel over time (*pseudocolor*), overlayed on greyscale images of vessels. Red scalebars show dilation of PA (*dark red*) and 1^st^ order capillary (*light red*). Astrocytic Ca2+ increase before and after dilation peaks in both PA and 1^st^ order capillary. Scalebar 20 µm. **H)** Mask of vessels (*purple*) and activity-dependent ROIs, detecting astrocytic responses prior to dilation peak (imaged marked with white star in H). **I)** Astrocytic Ca2+ averaged from all activity-dependent ROIs aligned to onset of all volitional whisking events of increasing duration. N= 20 FOV in 16 mice. **J)** The average max amplitude of astrocytic Ca2+ responses as a function of whisking duration. Pearson correlation R = 0.89, p = 0.016. N= 20 FOV in 16 mice. **K)** Average astrocytic Ca2+ activity in all ROIs during all volitional whisking events lasting >1s (*Green*) shows levels significantly above baseline both prior to and after PA dilation (*red*). Outline of 1 s.e.m. T-test vs baseline: Early Ca2+ max: p= 8.7*10-153 AUC: p=0.0033, Late Ca2+ max: p= 3.4*10-171, AUC p=0.00092, N=989 events, 20 FOV 16 mice. **L)** Ca2+ activity normalized to s.d. values for single whisking events lasting between 1½-2½ s, aligned to whisking onset (*yellow line*). <u>Above</u>: from astrocytic ROIs, <u>below</u>: from neuronal synaptic ROIs. **M)** Average 2-photon image from 105 µm below the cortical surface. *Red*: Texas red-dextran in vessel lumen, *Green*: GCaMP7 expressed in all neurons by the synapsin promotor. *Blue circle* outlines the synaptic ROIs within <15µm of the PA. Scalebar 20µm. **N)** Neuronal Ca2+ averaged from synaptic ROIs within <15µm of the PA, aligned to onset of all volitional whisking events of increasing duration. N= 19 FOV in 7 mice. **O)** The average max amplitude of neuronal Ca2+ responses from synaptic ROIs within <15µm of the PA as a function of whisking event duration. Pearson correlation R = 0.94, p = 0.0053. N= 19 FOV in 7 mice. **P)** Pupil imaged by an IR camera, used to detect the arousal state of the animal in relation to volitional whisking. **Q)** Average pupil size aligned to onset of all volitional whisking events of increasing duration. **R)** The average max amplitude of pupil dilation during volitional whisking of different durations. Pearson correlation R = 0.69, p = 0.12, N= 6 mice.

### Self-initiated whisking events trigger spatially recurrent astrocytic Ca2+ activities that rise before the dilations of the blood vessels and coincide with neuronal activation

To visualize astrocytic activity during the volitional NVC response, we used GCaMP6f virally introduced and regulated by the gfaABC1D promotor or a GMO strain constitutively expressing the fluorophore under the GLAST promotor. Given the current uncertainty with regard to astrocytic responses in the context of NVC, it was a strong priority that the intracellular Ca^2+^ changes were analyzed using an automated definition of the regions of activity. Automation minimizes the risk of bias associated with manually placing regions of interest (ROI). Hence, we first wrote a code that did automated event detection with a dynamic ROI definition similar to aQua(Wang, DelRosso et al. 2019, Krogsgaard, Sperling et al. 2023) (Supplemental figure S3). This method detected large and spatially heterogeneous dynamics in astrocyte Ca2+ activity (Supplemental Figure S4A). The average activity from these dynamic ROIs was correlated to whisking activity (Supplemental Figure S4B and C), and a small but significant response did occur (Supplemental Figure S4D). The dynamic ROI detection method often chose different regions between acquisitions, possibly because of a bias towards selecting regions with the highest level of Ca2+ activity. Such divergence was problematic as we wished to compare the same region between interventions. With that and since previous studies suggest that the larger astrocytic Ca2+ surges are not relevant for the recruitment of blood in neurovascular coupling (Institoris, Vandal et al. 2022), and indeed may reflect an integration of past events(Rupprecht, Duss et al. 2024), we decided to employ another automated ROI placement paradigm. Our alternative ROI definition was biased towards regions with spatially recurrent Ca2+ activity during NVC events triggered by volitional whisking (Figure 1G, Supplementary Figure S5). We found these by temporally aligning the images to the peaks of the whisking-evoked PA dilations (Figure 1G). Our algorithm then defined ROIs as the regions that, on average, had high levels of astrocyte activity preceding whisking-evoked dilation (Figure 1H, supplemental figure S5). These ROIs were then applied to detect Ca2+ activity in all acquisitions at their respective fields of view (FOVs) and normalized to the seconds prior to whisking initiation. We found the dilation-related astrocytic Ca2+ correlated well with whisking duration (Pearson correlation R = 0.894, p = 0.0164, N= 20 FOV in 16 mice, Figure 1I and J). The astrocytic Ca2+ activity peaked twice, both prior to and after the dilation (T-test comparing to baseline Early Ca2+ max: p= 8.7*10-153 AUC: p=0.0033, Late Ca2+ max: p= 3.4*10-171AUC p=0.00092, N=989, 21 FOV 16 mice) (Figure 1K). These two peaks were most prominent in the astrocytic ROIs closest to the PA (Supplemental Figure S6). Our astrocytic signal only reflected some degree of synchronicity with the whisking onset, as we found that the response to the individual whisking events showed high heterogeneity in timing, modality, and occurrence (Figure 1L, above).

To relate the astrocytic activity during volitional whisking to natural NVC, we also had to measure the neurons, as their responses are the basis of the energy demands that drive the vascular responses. To do so, we virally delivered the Ca2+ indicator GCaMP7 under the synapsin promotor in another mouse cohort (Figure 1M). Imaging of spontaneous neuronal Ca2+ activity in the syn-GcaMP7 mice showed high heterogeneity. Still, the average neuronal Ca2+ levels did change in correspondence with the volitional whisking events during rest and when the mouse was running. To localize the whisking-related signals, we identified active regions by correlating them to whisking onset (Supplemental Figure S7). This method revealed that most whisking-related neuronal activity appeared within ROIs with an area <1µm^2^ (Supplemental Figure S7). Given that murine synapses are around 1µm^2^, we propose that these are synaptic structures. ROIs larger than 1µm^2^ were considered dendrites, whereas somas were manually selected. Except for in somas and dendrites, the average activity in all neuronal compartments correlated well with the duration of the volitional whisking activity, (Supplemental Figure S7). In particular, the average response from synaptic ROIs in the proximity of the vessels (Pearson correlation R = 0.939, p = 0.00531, N=19 FOV in 7 mice Supplemental Figure S7 and Figure 1M-O). Similar to what was seen in the astrocytes, the synaptic-size neuronal activity during the individual volitional whisking events was also heterogeneous with regard to timing, modality, and occurrence (Figure 1L, below). The heterogeneity shows that whisking activity alone does not define the strength of the neuronal excitation. An explanation could be the different behavioral contexts of each volitional whisking.

One such modifying factor could be the activity of noradrenergic neurons during the whisking (Collins, Francis et al. 2023). To investigate the noradrenergic component in volitional NVC, we measured pupillary dilation as it is a straightforward, non-invasive method to quantify systemic NA release (Reitman, Tse et al. 2023) (Figure 1P and supplemental figure S8). To investigate how much arousal affected the astrocytic responses to self-initiated whisking, we aligned the changing pupil size to the onset of whisking events (Figure 1Q). There was indeed a change in the alertness and arousal state of the animal, even with the shortest whisking activities. The expected dilation was preceded by a constriction, likely because our definition of whisking onset required a half-second period without whisking (Figure 1Q). Pupil dynamics could thus be described as in sync with the whisking rather than as a response to the whisking. The size of the pupil dilations during whisking did not correlate well with whisking duration (Figure 1R, Pearson correlation R = 0.698, p = 0.1226, N=6 mice). Still, the dilations are significant and appear to be slightly larger to volitional whisking events longer than 1s. In accordance with increased activation of NA positive projections from the locus coeruleus during volitional whiskings >1s duration (Collins, Francis et al. 2023). We thus have some degree of brain state shift in the context of our volitional NVC events (Turner, Gheres et al. 2023).

### Stimulus strength during volitional whisking scales early astrocytic Ca2+ responses in spatial accordance with neuronal responses

Studies using anesthetized animals have shown that increasing the frequency of whisker stimulation triggers a more robust NVC response due to enhanced recruitment of the neurons in the barrel cortex (Jessen, Mathiesen et al. 2017). We could not control the frequency of the volitional whisking, so to increase sensory input, we placed a piece of sandpaper within reach of the whiskers (Figure 2A). Sandpaper stimulation resulted in larger PA dilations compared to whisking events without sandpaper (Figure 2B, Supplemental Figure S9B) (ANCOVA, p = 0.0065, N=20 FOVs in 16 mice). The sandpaper only influenced PA dilations during more extended whisking events (Supplemental Figure S9B). Thus, we focused our investigation on the cellular response to the enhanced sensory input on volitional whisking events longer than 1.5 seconds. Even though the sandpaper did not change the whisking behavior (Supplemental Figure S10), we reduced the variation of the response duration by excluding volitional whisking events that lasted >3 s. Measuring the PA expansion to 1.5-3 s of whisking in absolute values of µm^2^ confirmed the sandpaper effect (Figure 2C; Mixed-effect model, AUC: p=0.024, Max: p=0.0267, N= 20 FOVs in 16 mice). Interestingly, this effect was insignificant on the 1^st^ order capillaries (Figure 2D, Supplemental Figure S9C, Mixed-effect model, AUC: p=0.211, Max: p=0.129, N=10 FOVs in 7 mice). The difference suggests that the increased sensory input primarily strengthened blood flow recruitment in the deeper cortical layers but was too weak to trigger an additional blood flow response in more superficial layers. In the context of what is known about the cortical excitatory microcircuits, it agrees with an excitation of the neurons in layer IV before it reaches the Layer II neurons(Petersen 2019). Enlarging deflections with sandpaper did also not alter the neuronal responses overall in the FOV (Figure 2E, 2F and Supplemental Figure S9E). However, when we considered the synaptic ROIs in 15 µm circular bins from the PA, we found neuronal responses in synaptic ROIs within 30µm of the PA were substantially increased (Figure 2E, 2F and Supplemental Figure S9E, mixed model, Max early p=0.011, Max late p=0.0275, N=19 FOV, 7 mice). In accordance with this, we also found increased astrocytic Ca2+ responses in ROIs within a 30µm distance (Figure 2G, 2H and Supplemental Figure S9F and S9G, mixed model test, Max early=p0.00471, Max late p=0.0154 N=20 FOVs in 16 mice). Hence, early astrocytic activity was similarly affected and had the same spatial distribution as synaptic activity. This observation indicates that the first peak in the astrocytic response is due to the astrocytic proximity to the synapses, and that this astrocytic activity reflects the level of sensory-dependent neuronal excitation. Such spatial specificity is a requirement for astrocytic participation in the recruitment of the NVC response. The increased whisking deflection also significantly enlarged the second peak in the astrocytic Ca2+ response, which is proposed to prolong the blood flow response (Institoris, Vandal et al. 2022). This second increase exhibited less spatial specificity as it was also present in astrocytes >30µm from the PA (Figure 2G, 2H and Supplemental Figure S9F and S9G, mixed model test Max p=0.046 N= 20 FOVs in 16 mice).

**Figure 2.**
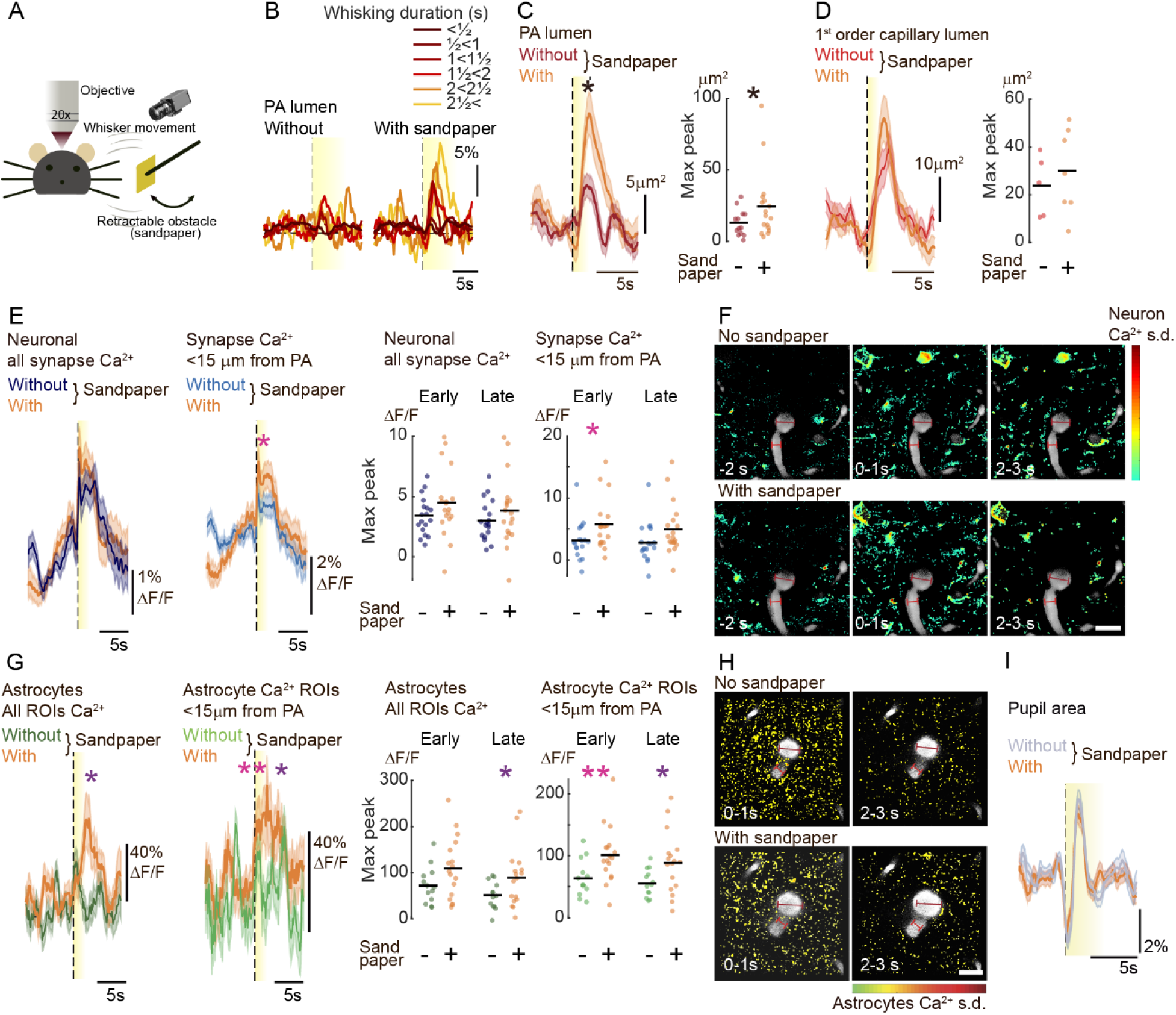
Increasing sensory input during volitional whisking elicited stronger vasodilations and increased Ca2+ activity in PA-proximal neuron and astrocyte structures. **A)** Cartoon of the setup showing the principle of the retractable obstacle. An infrared camera records whisker movements. A piece of sandpaper is repositioned by a motorized arm. **B)** Normalized average PA lumen during volitional whisking events of increasing duration of whisking, <u>Left</u>: without and <u>Right</u>: with sandpaper. **C)** Average PA dilation to all volitional whisking events 1½<3s long, measured in cross-area changes without (*dark red*) and with sandpaper (*orange*). Outline of 1 s.e.m., mixed model test, PA: AUC: p=0.024, Max: p=0.0267, N= 20 FOVs in 16 mice. **D)** Average 1^st^ order capillary dilation to all volitional whisking events 1½<3s long, measured in cross-area changes without (*red*) and with sandpaper (*orange*).Outline of 1 s.e.m., mixed model test, PA: AUC: p=0.211, Max: p=0.129, N= 10 FOVs in 7 mice. **E)** Average neuronal Ca2+ levels during all volitional whisking events 1½<3s long without (*dark blue*: All synaptic ROIs, *blue*: synaptic ROIs within <15µm from PA) and with sandpaper (*orange*). Outline of 1 s.e.m., Mixed model test: All: Max early p=0.088, Max late p=0.21, <15µm from PA: Max early p=0.021, Max late p=0.07, N= 19 FOVs in 7 mice. Scalebar 20µm. **F)** Averaged and normalized 2-photon images aligned to volitional whisking events from 1 s intervals, <u>above</u>: without and <u>below</u>: with sandpaper. Neuronal Ca2+ activity normalized to s.d. values per pixel over time (*pseudocolor*), before, at whisking onset, and 2. after. Overlayed *greyscale* image of vessels, red scalebar shows dilation of PA (*dark red*) and 1^st^ order capillary (*light red*). Scalebar 20µm. **G)** Average astrocytic Ca2+ levels during all volitional whisking events 1½<3s long without (*dark green*: All ROIs, *green*: ROIs within <15µm of PA) and with sandpaper (*orange*). Outline of 1 s.e.m., mixed model test, All: Max early p=0.077, Max late: p=0.049, 15µm from PA: Max early: p=0.00471, Max late p=0.0154 N=20 FOVs in 16 mice. **H)** Averaged and normalized 2-photon image aligned to volitional whisking events from 1 s intervals, <u>above</u>: without and <u>below</u>: with sandpaper. Astrocytic Ca2+ activity normalized to s.d. values per pixel over time (*pseudocolor*), at whisking onset, and 2. after. Overlayed vessels (*grey*); red, red scalebar shows dilation of PA (*dark red*) and 1^st^ order capillary (*light red*). The entire series is in Supplemental Figure S9G. Scalebar 20µm. **I)** Sandpaper did not affect the average pupil fluctuations during volitional whisking events 1½<3s long, without (*grey*) and with sandpaper (*orange*).

The modest and localized increase in neuronal activity achieved by sandpaper stimulation during volitional whisking suggests that the demand for increased blood flow is triggered by neuronal activity in deeper cortical layers and that these deeper layers relay to layer II/III neurons in close proximity to the PA. This neurovascular unit activity was not due to any arousing or aversive effect of the sandpaper, as the pupil expansion to volitional whisking events were similar with and without sandpaper (Figure 2I, Paired T-test, p = 0.531, N=6 mice). Thus, the effect of the sandpaper on neuronal and astrocytic activity, as well as the increased PA dilation, was best explained by additional sensory excitation produced by the larger deflections of the whiskers. Considering the spatial dependency and proportionality of astrocytic activity to neuronal responses, their early timing and position in relevant distance from the vascular inflow tract, we suggest this astrocytic activity could be directly involved in recruiting NVC responses to volitional whisking.

### During self-initiated whisking, early astrocytic Ca2+ activity facilitates blood flow recruitment in 1^st^ order capillaries in a NA-dependent manner

Astrocytic Ca2+ has been shown to depend strongly on NA levels (Paukert, Agarwal et al. 2014, Reitman, Tse et al. 2023), and so we hypothesized that astrocytes’ role in NVC could depend on cortical levels of NA. To test this, we targeted noradrenergic neurons in the LC using DSP-4. This neurotoxin eliminates NA-releasing neuronal projections from the LC, including to the barrel cortex (Iannitelli, Kelberman et al. 2023) and lowers cortical NA globally (Slezak, Kandler et al. 2019, Renden, Institoris et al. 2024). To confirm its effect on astrocyte Ca2+ activity, we measured the long-duration surges (> 5s long) with the dynamic ROI detection analysis (Supplemental Figure S3). We found, as expected, a significant drop in both the spatial (Figure 3A, T-test p=0.0043, N= 20 FOVs in 16mice) as well as temporal distributions of long-duration astrocytic Ca2+ events after DSP-4 treatment (Figure 3B, T-test p=0.0039, N= 20 FOVs in 16mice). We then evaluated pupil dynamics to confirm globally altered NA release during volitional whisking (Figure 3C). The effect of DSP-4 was a generalized pupil constriction, with baseline pupil diameter greatly reduced (Figure 3D, Paired T-test, p=4.2*10-5, N=6 mice) and a slight but non-significant reduction in whisking-induced dilation (Figure 3C, Paired T-test, p=0.224, N=6 mice). Remarkably, we observed a disruption in activity-dependent pupil synchronicity; pupils no longer first constricted and then dilated with the onset of volitional whisking (Figure 3E). Meanwhile, this depletion did not induce a change in whisking behavior (Supplemental figure S11).

**Figure 3.**
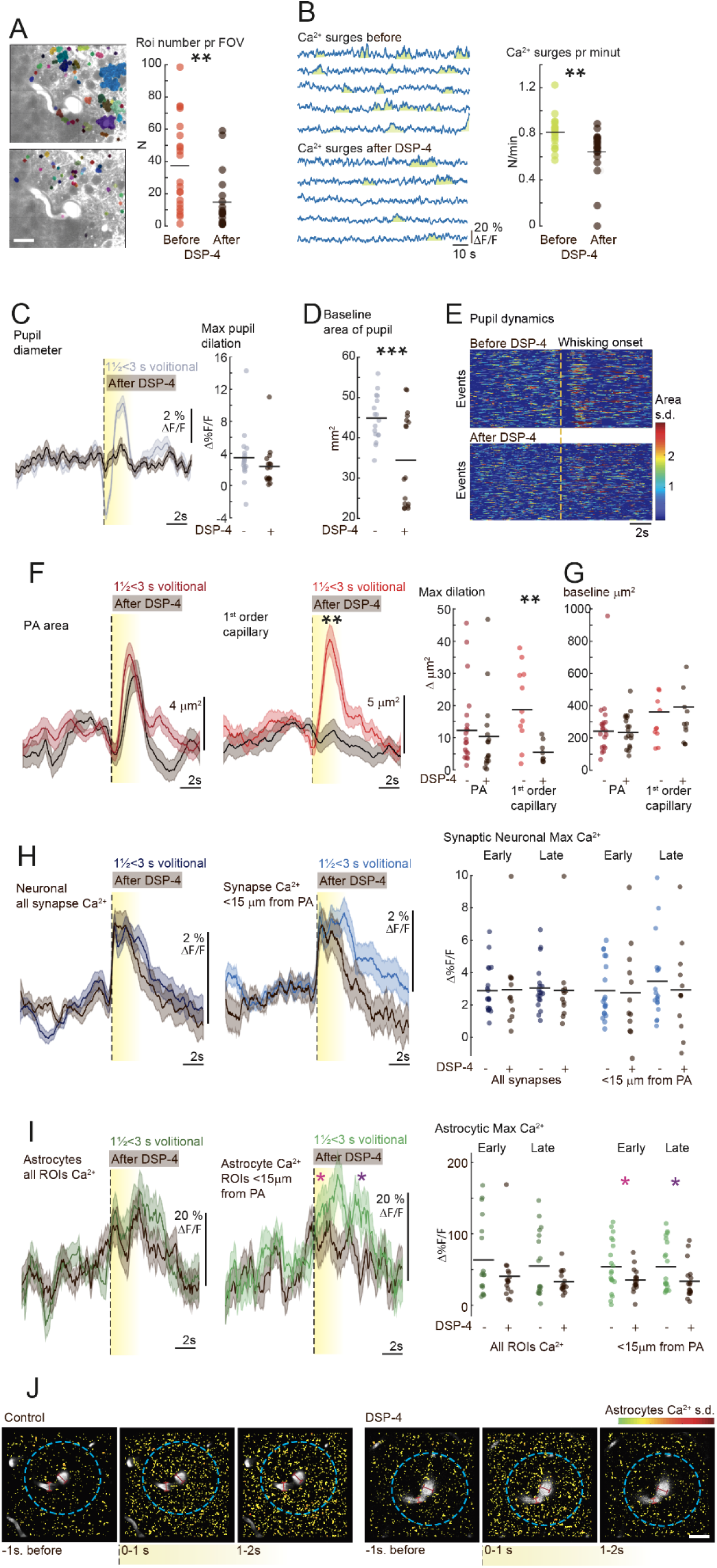
Locally disturbed astrocytic Ca2+ responses are accompanied by inhibited dilation of 1^st^ order capillaries during volitional NVC responses at rest after the pharmacological ablation of the LC. **A)** Fewer regions of dynamic astrocytic Ca2+ surges were detected following the locus coeruleus toxic DSP-4 administration. <u>Left</u>: Example of ROI detection results. <u>Right</u>: The average number of ROIs detected per FOV before (*red*) and after (*black*) DSP-4. T-test, p=0.0043, N=20 FOV per 16 mice. Scalebar 20 µm. **B)** <u>Left</u>: Example of long-duration (>5s) surges in astrocytic Ca2+ before (*above*) and after (*below*) DSP-4. <u>Right</u>: Quantified average occurrence of astrocytic Ca2+ surges from ROIs >30µm from the PA, before (*green*) and after DSP-4 (*black*). T-test, p=0.0039, N=20 FOV per 16 mice. **C)** <u>Left</u>: Despite DSP-4’s effect on the average pupil fluctuations during volitional whisking events 1½<3s long, <u>Right:</u> the maximal amplitude of the pupil dilations was not significantly different before (*grey*) or after DSP-4 (*black*). Paired T-test, p=0.224, N=6 mice. **D)** Baseline pupil area, averaged per mouse, was constricted after DSP-4 (*black*). Paired T-test, p=4.2*10-5, N=6 mice. **E)** Pupil area, normalized to standard deviation, during individual volitional whisking events lasting between 1½-3 seconds, illustrates a lack of synchronicity following DSP-4 treatment (*below*). **F)** Average vasodilations during volitional whisking events lasting between 1½-3 seconds, shaded outline is ± 1 s.e.m.. <u>Left:</u> PA dilation (*dark red*) did not change significantly after DSP-4 (*black*). <u>Middle:</u> 1^st^ order capillary dilation (*red*) was strongly reduced after DSP-4 (*black*). <u>Right:</u> Quantification of the average effect on vasodilations per FOV shows how the DSP-4 driven elimination of LC-based cortical NA release only affected 1^st^ order capillary dilations. Mixed model, PA: p=0.65 N=20 FOVs in 16 mice, 1^st^ order cap.: p=0.0045 N=10 FOVs in 7 mice. **G)** The effect of DSP-4 treatment was not due to an impact on the average baseline cross area of either PA (<u>left</u>) or 1^st^ order capillary (<u>right</u>). Mixed model, PA: p=0.579 N=20 FOVs in 16 mice, 1^st^ order cap.: p=0.578 N=10 FOVs in 7 mice. **H)** Average neuronal Ca2+ in synaptic ROIs during volitional whisking events lasting between 1½-3 seconds, shading is ± 1 s.e.m.. <u>Left:</u> Average from all ROIs (*dark blue*) did not change significantly after DSP-4 (*black*). <u>Middle:</u> Average of responses in synaptic ROIs within <15µm of PA (*blue*) were also not reduced by the DSP-4 (*black*). <u>Right:</u> Quantification of the average effect of the DSP-4 treatment on neuronal Ca2+ in ROIs per FOV. Mixed model, All ROIs: Early p=0.88. Late p=0.82, within 15µm of PA: early p=0.82, late p=0.57, N=19 FOV in 7 mice. **I)** Average astrocytic Ca2+ in ROIs during volitional whisking events lasting between 1½-3 seconds, shading is ± 1 s.e.m.. <u>Left:</u> Average from all ROIs (*dark green*) was not significantly changed after DSP-4 (*black*). <u>Middle:</u> Both early and late Ca2+ peaks in averages from ROIs within <15µm of PA (*green*) were significantly reduced by the DSP-4 (*black*). <u>Right:</u> Quantification of the average effect of the DSP-4 treatment on astrocytic Ca2+ in ROIs per FOV. Mixed model, All ROIs: Early Peak p=0.12, Late peak p=0.061 within 15µm of PA: Early peak p=0.0216 Late Peak: p=0.013, N=20 FOV in 16 mice. **J)** Averaged and normalized 2-photon images from volitional whisking events before (*left*) and after DSP-4 (*right*) from 1 s intervals. Ca2+ activity normalized to s.d. values per pixel over time (*pseudocolor*), before, at and 1 sec after whisking onset. Overlayed vessels (*grey*); red scalebar shows max dilation of PA (*dark red*) and 1^st^ order capillary (*light red*); *dashed blue line* shows a 30µm distance from PA. Scalebar 20 µm.

We then investigated whether the volitional NVC response remained intact after the disruption in the NA release. While PAs retained appropriate dilation amplitudes, we found DSP-4 treatment had abolished dilatory activity in 1^st^ order capillaries (Figure 3F, mixed model, PA p=0.65, N= 20 FOVs in 16mice, 1^st^ order cap.: p= 0.0045, N= 10 FOV in 7 mice). As NA can have a vasoconstricting effect (Montalant, Kiehn et al. 2024), we wondered whether the depletion of cortical NA levels had established a higher baseline lumen-area in the vessels. This was not the case, as neither the PA nor the 1^st^ order capillary was expanded in the DSP-4 treated animals (Figure 3G, Mixed model test, PA p=0.70, N= 20 FOVs in 16 mice, 1^st^ order cap.: p= 0.88, N= 10 FOV in 7 mice). We then evaluated whether a reduced local neuronal activity could explain the reduced 1^st^ order capillary dilation response. We found that the neuronal excitation by volitional whisking was not decreased after the DSP-4 treatment (Figure 3H, Supplemental Figure S12A and B, Mixed model, All ROIs: p=0.88, within 15µm of PA: p=0.82, N=19 FOV in 7 mice). Now, the question was whether the attenuation of the larger astrocytic Ca2+ activities after the reduction in NA levels had altered the early and local astrocytic responses to volitional whisking. Surprisingly, the effect on the astrocytic responses to volitional whisking was not strong, and the early Ca2+ responses were generally intact. Still, we did find a slight yet significant reduction in the ROIs bordering the vascular inflow tract (Figure 3I and 3J, Supplemental Figure S12C-E, mixed model test, p=0.0216, N= 20 FOVs in 16 mice). The general integrity of the signal implies that astrocytes still responded to synaptic activity but that NA depletion had blunted a part of their response. The post-dilation peak in astrocyte Ca2+ was also significantly attenuated in vessel-adjacent ROIs (Figure 3I and 3J, Supplemental Figure S12C-E, mixed model test, within 15µm of PA: p=0.012, All ROIs: p=0.061 N= 20 FOVs in 16mice).

These results indicate that the early astrocytic Ca2+ activity in cells bordering the inflow tract plays an essential role in the local regulations of the vascular tone of 1^st^ order capillaries during volitional whisking. Furthermore, this precise spatial and temporal astrocytic regulation is dependent on intact cortical NA signaling.

### Self-initiated whisker movements show earlier neuronal Ca2+ activity than evoked deflections, while astrocytic Ca2+ and vascular responses are unchanged

As the sensory inflow in self-initiated whisking can be preceded by activity in several neuronal networks(Eggermann, Kremer et al. 2014, Kinnischtzke, Simons et al. 2014), we wondered whether the cellular processes leading to a NVC response were proactive. To test this, we compared the NVC response in volitional whisking to an evoked air puff stimulus (Figure 4A). While natural whisking is an active and expected deflection, the air puff stimulation represents an unexpected sensory event. To avoid arousal or startle response, we minimized air pressure to cause the smallest visible deflection and ensured that the air puff only hit the whiskers. We also thoroughly trained the animal to air-puff stimulation before the experiments. Even so, air puff stimulation made the animal run in approximately 17% of the trials, suggesting some startling effect. These cases were discarded, and only stimulations that did not result in running were included.

**Figure 4.**
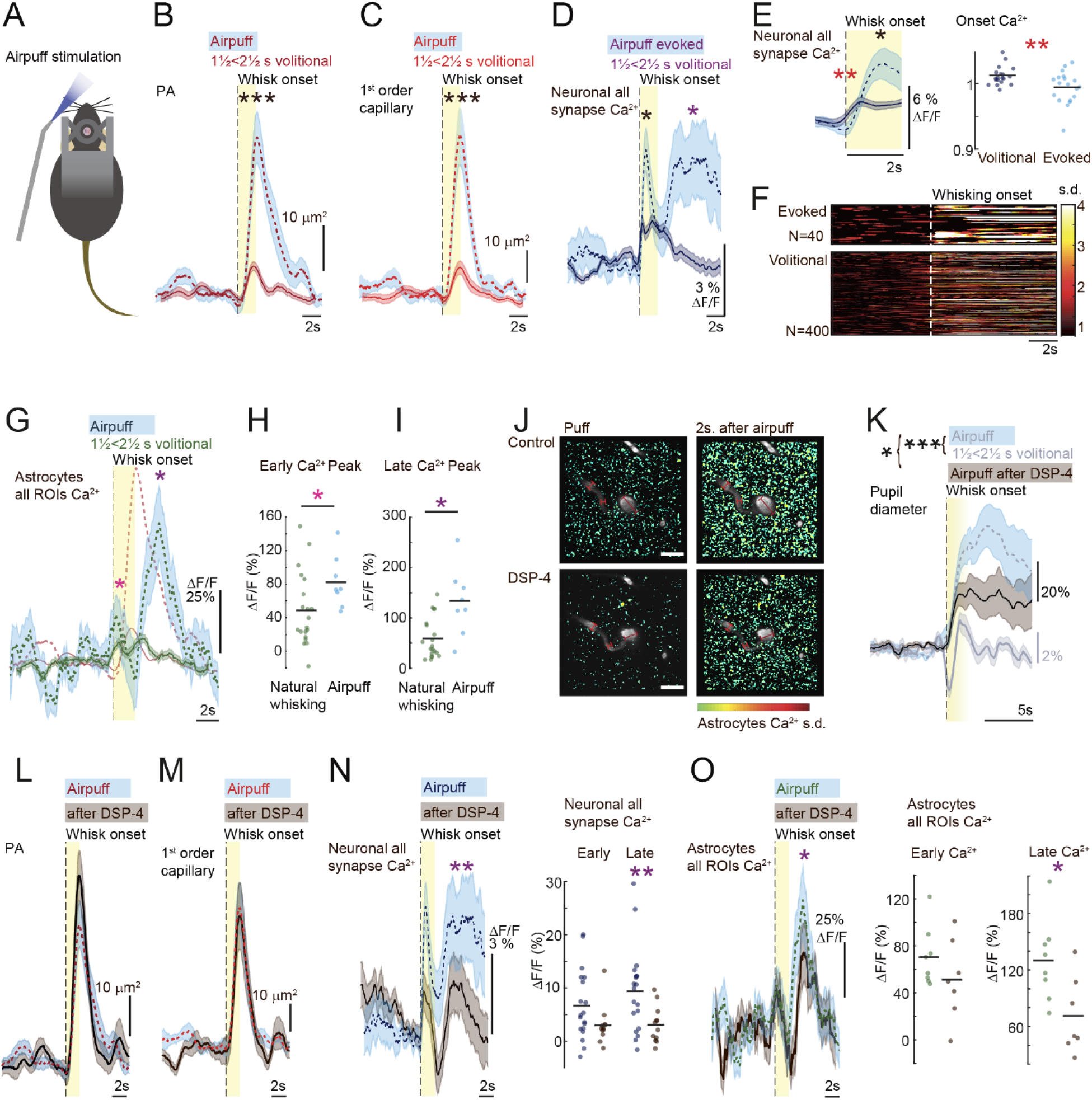
Evoked NVC response did not depend on cortical noradrenergic levels, but astrocytic and vasoactive responses were otherwise comparable to volitional whisking. **A)** Cartoon of the evoked whisker stimulation using air puff **B)** Average PA vasodilation during volitional whisking events lasting between 1½-2½ seconds (*dark red, solid line*) was significantly smaller than during 2s air-puff evoked whisking (*dashed line*, ± 1 s.e.m.. outlined in *light blue*). Welch’s t-test, p=0.014, Air puff N=8 FOVs in 6 mice vs natural whisking N= 20 FOV in 16 mice, 1^st^ order cap: p=0.0126, Air puff N=8 FOVs in 6 mice vs natural whisking N= 10 FOV in 7 mice, PA: p=0.65 N=20 FOVs in 16 mice, 1^st^ order cap.: p=0.0045 N=10 FOVs in 7 mice. **C)** Average 1^st^ order capillary vasodilation during volitional whisking events lasting between 1½-2½ seconds (*red solid line*) was significantly smaller than during 2s air-puff evoked whisking (*dashed line*, 1 s.e.m. outline in *light blue*). Welch’s t-test, p=0.0126, Air puff N=8 FOVs in 6 mice vs natural whisking N= 10 FOV in 7 mice. **D)** Average neuronal Ca2+ during volitional whisking events lasting between 1½-2½ seconds (*dark blue solid line*) were significantly smaller than during 2s air-puff evoked whisking (*dashed line*, 1 s.e.m. outline in *light blue*). Paired t-test, Max early peak (*black*) p = 0.013, Max late peak (*dark blue*) p=0.0194, N=19 FOV in 7 mice, **E)** Left, Average neuronal Ca2+ from synaptic ROIs zoom in from C). Right: Quantified onset Ca2+ levels (0-100ms after whisking onset) averaged pr FOV. Neuronal Ca2+ in all synaptic ROIs rose significantly earlier to volitional whisking (*dark blue solid line*) than to evoked whisking (*dashed line*, 1 s.e.m. outline in *light blue*). Paired t-test, Onset level (*red stars*) p=0.0089 N=19 FOV in 7 mice, *black star* same as in D). **F)** Neuronal Ca2+ activity average from all synaptic ROIs normalized to s.d. values (*pseudocolor*). For single whisking events, aligned to whisking onset (*yellow line*). <u>Above</u>: Air-puff evoked, <u>Below</u>: from volitional whisking events lasting between 1½-2½ s. **G)** Astrocytic average Ca2+ from all ROIs during volitional whisking events lasting between 1½-2½ seconds (*dark green solid line*) was significantly smaller than during 2s air-puff evoked whisking (*dashed line*, 1 s.e.m. outline in *light blue*). Average PA dilations from B) (*dark red dashed line*) reveal how both responses rose in two peaks, one before and one after the vasodilations. **H)** Peak amplitude in the “early” Ca2+ response from G) shows that evoked responses were larger than the volitional (*pink star*). Welch’s t-test, Early peak Max p = 0. 032 puff N= 8 FOVs in 6 mice vs. natural whisking N= 20 FOV in 16 mice. **I)** Peak amplitude of the late response in the Ca2+ response peak from G) was larger in the evoked responses than the volitional (*purple star*). Welch’s t-test, Late peak Max p=0.018 Air puff N= 8 FOVs in 6 mice vs natural whisking N= 20 FOV in 16 mice. **J)** Average 2-photon image from air puff evoked events before (*above*) and following DSP-4 (*below*). Ca2+ activity normalized to s.d. values per pixel over time (*pseudocolor*), 2 and 4 seconds after air puff. Overlayed vessels (*grey*); red, red scalebar shows dilation of PA (*dark red*) and 1^st^ order capillary (*light red*). Scalebar 20µm. **K)** Average pupil dilations during volitional whisking events lasting between 1½-2½ seconds (*grey solid line*) were significantly smaller than during 2s air-puff evoked whisking (*black dashed line*, 1 s.e.m. outline in *light blue*). Paired t-test, p=0.000128, N=6 mice. DSP-4 reduced the response (*black solid line*, s.e.m. outline in *dark grey*), paired t-test p=0.014, 6 mice. **L)** Average PA vasodilations during 2s air-puff evoked whisking (*Dark red dashed line*, 1 s.e.m. outline in *light blue*) were not significantly changed by DSP-4 black (*solid line, black*, s.e.m. outline in *dark grey*). Paired t-test, p=0.718, N=8 FOV in 6 mice. **M)** Average 1^st^ order capillary vasodilations during 2s air-puff evoked whisking (*Red dashed line*, 1 s.e.m. outline in *light blue*) were not significantly changed by DSP-4 (*black solid line*, s.e.m. outline in *dark grey*). Paired t-test, p=0.632 N=8 FOV in 6 mice. **N)** Neuronal Ca2+ averaged from all synaptic ROIs during 2s air-puff evoked whisking. (All ROIS, *Dark blue dashed line*, 1 s.e.m. outline in *light blue*), was significantly reduced by DSP-4 (*black solid line,* s.e.m. outline in *dark grey*). Paired t-test, All ROIs Early peak p=0.052, Late peak p= 0.0053, ROIs<15µm Early peak p=0.10, Late peak p= 0.26, N=19 FOV in 7 mice. **O)** Astrocytic Ca2+ averaged from all ROIs during 2s air-puff evoked whisking (*Dark green dashed line*, 1 s.e.m. outline in *light blue*) was only significantly changed by DSP-4 (*black solid line*, s.e.m. outline in *dark grey*) in the late peak of the response. Paired t-test, Early peak p=0.24, Late peak p=0.018, N=8 FOV in 6 mice.

Air puff stimulation resulted in robust vasodilations of both the PA and the 1^st^ order capillary. These were larger than the dilations during self-initiated whisking events of similar duration (Figure 4B and 4C, Welch’s t-test, PA: p=0.014, Air puff N=8 FOVs in 6 mice vs. natural whisking N= 20 FOV in 16 mice, 1^st^ order cap: p=0.0126, Air puff N=8 FOVs in 6 mice vs natural whisking N= 10 FOV in 7 mice). The increase in both PA and 1^st^ order capillary distension was matched by increased neuronal Ca2+ activity (Figure 4D, Welch t-test, Max early peak p = 0.013, Max late peak p=0.0194 N=19 FOV in 7 mice) across all synaptic ROIs, independent of distance from the blood vessel. Such an increase in neuronal excitation justifies the substantial blood flow response. In addition, the different initiation pathways recruited in evoked versus volitional whisker deflection allow for the latter to involve a preemptive excitatory signal to condition the somatosensory cortex for an anticipated sensory input. Indeed, the neuronal response to volitional whisking was slightly earlier than to evoked whisker deflections (Figure 4E and F, paired t-test, Onset p=0.0089 N=19 FOV in 7 mice). However, this preemptive signal did not appear to directly affect the timing of the blood flow recruitment, as the dilation onset of both PAs and 1^st^ order capillaries was similar between evoked and volitional events (Figure 4B and 4C).

As early astrocytic Ca2+ activity has often been questioned in evoked sensory responses, we wondered if we had observed an astrocytic activity specific to self-initiated NVC responses. However, when we investigated the astrocytic Ca2+ responses within the ROIs identified from the volitional whisking we again found that astrocyte activity rose both prior to and after the peak vasodilation (Figure 4G). Comparing the amplitude of these two Ca2+ transients to their volitional counterparts, the early astrocytic Ca2+ response was slightly but significantly larger than in self-initiated NVC (Figure 4G and 4H). In contrast, the second and delayed activity was substantially larger in the air puff condition (Figure 4G and 4I, Welch’s t-test, Early peak Max p = 0. 032, Late peak Max p=0.018 Air puff N= 8 FOVs in 6 mice vs. natural whisking N= 20 FOV in 16 mice). Thus, we found no difference in astrocytic Ca2+ activity between evoked and volitional NVC responses that could not be explained by heightened neuronal activity. The increased early astrocytic Ca2+ peak matched the increased synaptic activity, and the much larger second peak the size of the evoked dilation. In summary, the more prominent activation of neurons, astrocytes, and vasculature in the evoked as compared to volitional whisking suggests that the comparably bigger whisker deflection of the air puff stimulation led to a stronger sensory input and more widespread excitation (Figure 4B-J).

### Vascular responses to evoked whisking events did not depend on cortical NA levels or reduced astrocytic Ca2+ surge activity, but neuronal responses did

One possible confounding factor to this assumption was that the pupil dilated much more during air-puff evoked than volitional whisking (Figure 4K, Paired t-test, p=0.000128, N=6 mice). Thus, we suspected some arousal effect of the air-puff stimulation despite our precautions to avoid startling the animal. A small arousal response would induce NA release in the cortex, which could influence the neuronal and astrocytic responses to the puff-evoked whisking. Hence, we evaluated the response to the evoked deflections in the NA-depleted mice. While DSP-4 slightly decreased pupil dilations after air puff stimulation (Figure 4K, Paired T-test, p=0.014, N=6 mice), the vascular response in both the PA and 1^st^ order capillaries was unchanged (Figure 4L and 4M, Supplemental figure S13B; t-test, PA: p=0.718, 1^st^ order cap.: p=0.632 N=8 FOV in 6 mice).

Meanwhile, we found that the neuronal response was reduced. This reduction was primarily within the second, later peak and in synaptic ROIs more than 30µm distance from the vessel (Figure 4N; t-test, All ROIs, early peak: p=0.052, Late peak: p=0.0053 N=19 FOV in 7 mice Supplemental figure S13C and S13D). In accordance with the unchanged NVC response, the synaptic responses within 30µm were unaffected by reduced NA (Supplemental figure S13C and S13D). Likewise, DSP-4 did not alter the early, pre-dilation astrocytic Ca2+ response (Figure 4O, Supplemental figure S13E-G, t-test p=0.126, N=8 FOV in 6 mice). In contrast, the late, post-dilation astrocytic Ca2+ response to air puff was decreased (t-test p=0.018, N=8 FOV in 6 mice, Figure 4P, Supplemental figure S13E-G). The lack of effect from the DSP-4 treatment indicated that NA did not substantially contribute to air puff-induced NVC response. However, it was remarkable that neuronal response was lowered without a similar reduction in vascular responses. Another striking difference between the self-initiated and evoked NVC responses was that the DSP-4 attenuation of both cortical NA levels and astrocytic Ca2+ surges neither altered early astrocytic Ca2+ response nor reduced 1^st^ order capillary dilation. This suggests a substantial difference in the blood flow recruiting mechanisms between evoked and volitional NVC responses.

### During locomotion, the localized whisking-based NVC response no longer matched neuronal or astrocytic activity

So far, we have only considered NVC responses to whisker deflections in resting animals. Whisking, however, plays an essential role in locomotion, as mice continuously whisk while moving (Supplemental Figure S2). In addition, any locomotion event would be preceded by a period of volitional whisking before the onset of locomotion. This initial whisking is a reasonable surveillance behavior prior to an increase in movement speed, as rodents primarily rely on their whiskers to explore the surroundings. Evidently, the excessive whisking during locomotion should provide a strong excitation of the sensory pathway. Locomotion is also associated with arousal, which entails the release of NA to many parts of the sensory cortices, including the barrel cortex (Shimaoka, Harris et al. 2018). As astrocytes are known to respond to NA during arousal, one could think that the widespread NA release from the LC modified astrocytic contributions to volitional NVC. As observed in previous studies (Ding, O’Donnell et al. 2013, Tran, Peringod et al. 2018, Oe, Wang et al. 2020, Renden, Institoris et al. 2024), we noted that locomotion induced both robust vasodilations and large, synchronous, and widespread astrocytic Ca^2+^ responses across the entire field of view (Figure 5A, 5B and 5C). The locomotion-related astrocytic Ca2+ peaked later than the dilation of both the PA and the 1^st^ order capillary (Supplemental Fig S14B, Wilcoxon test, PA p = 3.33e-16, 1^st^ order capillary p = 3.8e-11 N=177 locomotion events in 16 mice). More relevant to the recruitment of blood flow, the onset of the astrocytic Ca2+ response preceded vasodilation onset (Figure 5D, Wilcoxon test, p = 0.047, N=177 locomotion events in 16 mice). In addition, both astrocytic and vascular responses began after whisking onset and prior to locomotion (Figure 5D, Mann-Whitney, ast. Ca^2+^ p = 5.9*10-5, PA p = 0.0029, N=177 Events in 16 mice). That is, locomotion-related changes in blood flow and cellular activity in the barrel cortex occurred well before the animal began to move. This implies that the initial response in the barrel cortex is primarily due to the increased sensory input as a consequence of the whisking. As expected, we found a remarkable pupil expansion compared to whisking events of similar duration in the resting animals (Figure 5E, paired t-test, p=5,9*10-6, N=6 mice). Revealing a more substantial brain state switch and cortical release of NA upon initiation of the locomotion.

**Figure 5.**
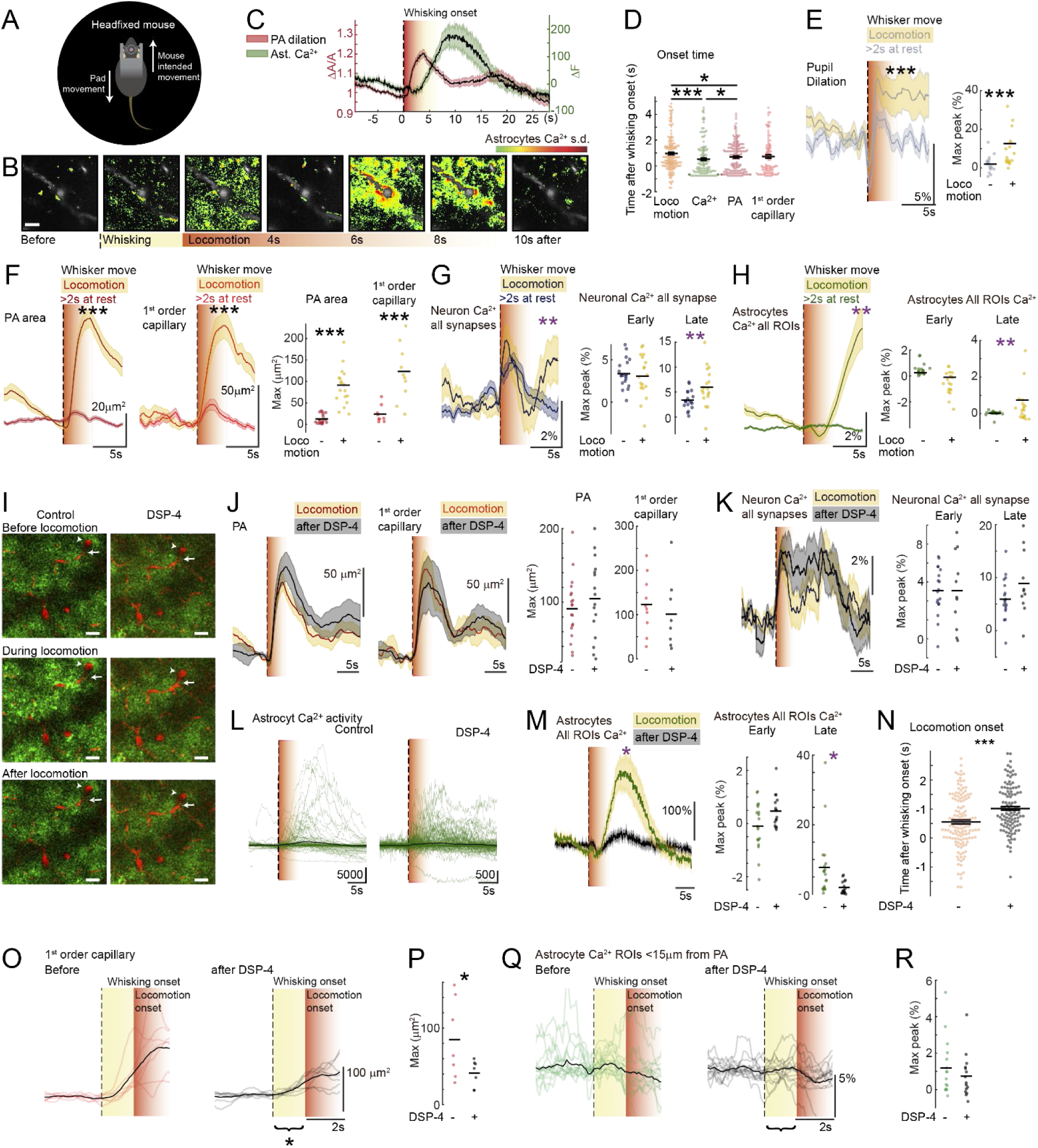
During locomotion, the localized whisking-based NVC response no longer matched neuronal or astrocytic activity, and none of the early responses were dependent on locus coeruleus release of NA. Still, the resting period before locomotion was prolonged, as was the recruitment of the 1^st^ order capillary. **A)** Cartoon of Neurotar system recording speed and direction of movement. **B)** Averaged images in 1s intervals before and during locomotion. Vessel lumen (*grey*) and astrocytic Ca2+ levels in s.d. normalized per pixel over time (*pseudocolor*). Scalebar 20µm. **C)** Averages of all locomotion events aligned to whisking onset. The orange area marks the period after whisking onset wherein locomotion began. PA vasodilation and Ca^2+^ activity in all astrocytic ROIs, N=177 locomotion events in 16 mice. **D)** Onset time of activity from onset of whisking associated with locomotion. *Orange*: Locomotion; *green*: astrocyte Ca2+; *red*: PA dilation; *light red*: 1^st^ order capillary dilation. The Ca^2+^ response (Wilcoxon test, p = 5.9*10-5) and the vasodilation (Wilcoxon test, p = 0.029) started before the locomotion while all responses lagged behind the whisking onset. The onset of astrocytic Ca2+ responses was significantly earlier than the vasodilation (Wilcoxon test, p = 0.047). N=177 locomotion events in 16 mice. **E)** <u>Left</u>: Average pupil fluctuations during volitional whisking events >2s long (*grey, solid line*) were significantly smaller than during >2s locomotion (1 s.e.m. outline in *yellow*). <u>Right</u>: Quantification of max amplitudes averaged pr mouse, Paired t-test p = 5.9*10-6, N=6 mice. **F)** Vasodilations were significantly larger during >2s long whisking events in locomotion than in rest. <u>Left</u>: Average PA dilations (s.e.m. outline *dark red*) during rest (*dark red outline*) and locomotion (s.e.m. outline *yellow*). <u>Middle</u>: 1^st^ order capillary dilation (*red*) during rest (s.e.m. outline *red*) and locomotion (s.e.m. outline in *yellow*). <u>Right</u>: Quantification of max amplitudes averaged pr. FOV, Mixed model: PA p = 4.1-10-10 N=20 FOVs in 16 mice, 1^st^ order cap.: p=9.4*10-5 N=10 FOVs in 7 mice. **G)** <u>Left</u>: Average neuronal Ca2+ from synaptic ROIs (*dark blue*) were significantly larger in >2s long whisking events during locomotion (s.e.m. outline *yellow*) than in rest (s.e.m. outline *dark blue*). <u>Right</u>: Quantification of max amplitudes averaged pr FOV, Mixed model test: Early peak: p=0.68, Late peak p = 0.0060 N=19 FOVs in 7 mice. **H)** <u>Left</u>: Astrocytic Ca2+ averaged from all ROIs (*dark green*) were significantly larger in >2s long whisking events during locomotion (s.e.m. outline *yellow*) than in rest (s.e.m. outline *dark green*). <u>Right</u>: Quantification of max amplitudes averaged pr FOV, Mixed model: Early peak: p = 0.28, Late peak: p=0.0016, N=20 FOVs in 16 mice. **I)** Example fluorescence images before (top), during (middle), and after (below) locomotion before (left) and after (right) administration of DSP-4. Arrows without stalk point to a penetrating arteriole and with stalk point to a 1^st^ order capillary. Scalebar 100µm. **J)** DSP-4 did not significantly affect the average vasodilations during >2s locomotion events. <u>Left</u>: Average PA dilations before (*dark red, yellow* s.e.m. outline) and after DSP-4 (*black, grey* s.e.m. outline). <u>Middle</u>: 1^st^ order capillary dilation (*red, yellow* s.e.m. outline) and after DSP-4 (*black, grey* s.e.m. outline). <u>Right</u>: Quantification of max amplitude per locomotion event. Mixed model: PA p=0.39, control N=20 FOV 16 mice and DSP-4 N=19 FOV in 14 mice, 1^st^ order capillary: p=0.49, control N= 10 FOV and DSP-4 N=11 FOV in 7 mice. **K)** <u>Left</u>: Average neuronal Ca2+ in all synaptic ROIS during locomotion (*dark blue, yellow* s.e.m. outline) was slightly increased after DSP-4 (*black, grey* s.e.m. outline). <u>Right</u>: Quantification of max amplitudes per FOV. Mixed model: Early peak: p = 0.68, Late peak: p=0.12 control N=19 FOV in 7 mice. **L)** Astrocytic Ca2+ dependent fluorescence averaged over all ROIS during individual locomotion events before (left) and after DSP-4 (right). **M)** <u>Left</u>: Average astrocytic Ca2+ in all ROIS during locomotion before (*dark green, yellow* s.e.m. outline) and after DSP-4 (*black, grey* s.e.m. outline). The late peak in astrocytic Ca2+ was reduced after DSP-4. <u>Right</u>: Quantification of max amplitudes per FOV. Mixed model, Early peak: p=0.22, Late Ca: p=0.017, control N=20 FOV in 16 mice, DSP-4 N=19 FOV in 14 mice. **N)** The delay from whisking onset to locomotion onset in all locomotion events was shorter before (*orange)* than after DSP-4 treatment (*black*). Mean with C.L. are shown. Mann-Whitney test p=3.94*10-5, Control: N=177 events in 16 mice and DSP-4 N=142 locomotion events in 13 mice. **O)** Dilation responses in 1^st^ order capillaries in >1.5 s whisking (*yellow*) leading up to locomotion (*red*) averaged pr FOV before (<u>left</u>, *red traces*) and after DSP-4 treatment (<u>right</u>, *black traces*). **P)** Mean max peak pr FOV. Mann-Whitney test, p=0.042, control: N=8 FOV in 7 mice, after DSP-4 N=8 FOV in 6 mice. **Q)** Astrocytic Ca2+ response from ROIs <15µm from PA in >1.5 s whisking (*yellow*) leading up to locomotion (*red*) averaged pr FOV before (<u>left</u>, *green traces*) and after DSP-4 treatment (<u>right</u>, *black traces*). **R)** Mean max peak pr FOV. Mann-Whitney test, p=0.36 Astrocytic Ca2+ response from ROIs <15µm from PA, control: N=8 FOV, in 7 mice, after DSP-4 N=8 FOV in 6 mice.

To investigate the contribution of this locomotion-related brain state switch to the whisking-based NVC response, we compared it to responses during self-initiated whisking events of similar duration in the resting animal. As expected, both PA and 1^st^ order capillary dilations were substantially increased by locomotion (Figure 5F, mixed model test, PA: p=4,1*10-10, N= 20 FOV in 16 mice, 1^st^ order cap: p=9,3*10-5 N= 10 FOV in 7 mice). Synaptic activation did not explain this augmentation, as neuronal Ca2+ was not concurrently increased. Meanwhile, a second Ca2+ peak was formed in the neuronal response, with levels significantly above what was seen in the resting animal (Figure 5G, mixed model, Early peak: p=0.67, Late peak: p=0.0061 N=19 FOV in 7 mice, supplemental Fig S14C and D). This second peak likely reflects the prolonged whisking during the locomotion. This second peak cannot be the recruiting factor in the enlarged vascular response but may partly explain the prolongation (supplemental Figure S14D). Considering the astrocytic Ca2+ responses, activity also increased during locomotion, and this increase was also only significant in the post-dilation part of the response (Figure 5H, mixed model test, Early Ca: p=0.26, Late Ca: p=0.0016, N=20, FOV, 16 mice, supplemental Fig S14E). As with the neuronal synaptic activity, the second and more prominent peak in astrocytic Ca2+ seems to rise as the PA begins to constrict back to its pre-whisking size (Figure 5C). Thus, the excess vasodilation during locomotion was not explained by increased activity in the sub-cellular neuronal and astrocytic structures included in this study. As these were selected because of their activation in volitional and evoked whisking during rest, it points to a very different activity pattern during locomotion.

### Depletion of locus coeruleus-dependent NA release did not affect neurovascular coupling during locomotion but delayed the transition from resting whisking to locomotion and the recruitment of the 1^st^ order capillary

To investigate this further, we returned to the DSP-4 treated animals. In these, we found that the vasodilatory response to locomotion was not affected (Figure 5I and 5J, Mixed model, PA: p=0.39, control N=20 FOV 16 mice and DSP-4 N=19 FOV in 14 mice, 1^st^ order cap: p=0.49, control N= 10 FOV and DSP-4 N=11 FOV in 7 mice). Similarly, there was no significant change to neuronal Ca2+ responses following the DSP-4 treatment (Figure 5K, Supplemental Figure S14F, mixed model test, Early peak: p=0.68, late peak p=0.12, N=19 FOV in 7 mice). Running-evoked astrocytic Ca^2+^ activity has previously been shown to be due to noradrenaline release from the locus coeruleus (Slezak, Kandler et al. 2019). Therefore, we expected the astrocytic Ca2+ responses to decrease in the NA-depleted animals. Indeed, after the administration of DSP-4, the bigger second peak of the astrocytic Ca^2+^ response was almost completely abolished (Figure 5L). However, looking closer at the remaining astrocytic Ca2+ dynamics after the DSP-4 treatment, the early response appeared to be intact (Figure 5L, right). This early activity was easy to overlook prior to the DSP-4 treatment, as the second, NA-based Ca2+ peak ramps-up upon locomotion initiation. We quantified the early astrocytic Ca2+ response and found that it was not reduced after the DSP-4 treatment (Figure 5M, Supplemental Figure S14G, mixed model test, Early Ca: p=0.22, Late Ca: p=0.017, control N=20 FOV in 16 mice, DSP-4 N=19 FOV in 14 mice). This means that the early peak in astrocytic Ca2+ still correlates with neuronal activity during locomotion and does not disappear with reduced cortical levels of NA. In our locomotion investigation, we aligned the cellular and vascular activity to the whisking onset rather than the locomotion onset. We did this because this study was done in the whisker barrel cortex, where the main excitatory input is expected to be from whisker deflections. Interestingly, the only behavior change we could detect following DSP-4 treatment was with regard to the interval between whisking onset and locomotion onset (Figure 5N, Supplemental Figure S11). Untreated animals would only whisk 0.55±0.14s before transitioning to the subsequent behavior (Figure 5N, Mann-Whitney test p=3.94*10-5, Control: N=177 events in 16 mice and DSP-4 N=142 locomotion events in 13 mice). Following depletion of the LC projections, the mouse whisked for 1±0.13s before locomotion onset. We wondered whether this could be related to the impaired vasodilations in 1^st^ order capillary we had found during volitional whisking in the resting animal (Figure 3). We evaluated whether a similar blockade was present before locomotion, which proved difficult as we could only focus on locomotion events preceded by a substantial period of volitional whisking. Thus, our final investigation only included locomotion events with a >1.5s whisking period before locomotion onset. Despite excluding many locomotion events, we could indeed detect a significant reduction in the initial dilation of the 1^st^ order capillary (Figure 5O-P, Mixed model test, p=0.042 control: N=8 FOV, in 7 mice, after DSP-4 N=8 FOV in 6 mice). Confirming that local blood flow regulation was impaired until the mice transitioned into the locomotion-related brain state. We thus also expected a similar effect from DSP-4 on astrocytic Ca2+ responses. We focused on our previously identified early peak in astrocytic activity in vessel-adjacent ROIs and compared the maximal Ca2+ level before and after DSP-4 within the first second in whisking-before-locomotion events (Figure 5Q-R, Mixed model test, p=0.36 control: N=8 FOV, in 7 mice, after DSP-4 N=8 FOV in 6 mice). While we did find a reduction, the variance in astrocytic Ca2+ data meant we could not conclude that this reduction in the early astrocytic response prior to locomotion was significant. Hence, it is still possible that the decline in astrocytic Ca2+ surges is the critical component in recruiting 1^st^ order capillaries in the NVC response during this behavioral transition (Figure 3A-B). The behavioral disturbance elicited by DSP-4 treatment could suggest an important regulatory role for NA-dependent astrocytic Ca2+ levels, enabling sufficient localized blood flow in the cortical processing of sensory information.

## Discussion

This study is the first investigation of the astrocytic contribution to NVC response to self-initiated whisking. To quantify this, we developed a method that identified the location of astrocyte activities with spatial and temporal recurrency in relation to the NVC event. We found astrocytic activation within these regions during all NVC events, regardless of whether the increased sensory input was volitional or evoked, during rest or locomotion. Using this analysis tool, we show that astrocytes may partake in blood flow recruitment during any NVC response in the whisker barrel cortex and during any change in behavior associated with increased whisker deflection. Despite this, we found that the recruitment of blood flow is not similarly dependent on astrocyte activity in different behavioral contexts. The difference lies in whether the animal is taking or receiving a sensory stimulus and whether it is running or not. When the sensory inflow was self-initiated, the astrocytic response could be attenuated by a reduction of cortical noradrenaline levels. This reduction impaired the recruitment of the 1^st^ order capillary and, thus, the fine-tuned regulation of local blood flow. The disruption of the neurovascular coupling might even impair sensory processing as the mouse now whisked for a longer time before initiating a locomotion. The NVC response to evoked whisking and during locomotion was not reduced. Neither was the early astrocytic peak reduced in these events, so our data cannot exclude an astrocytic contribution to NVC in evoked whisking or during locomotion. Our results suggest that astrocytic contribution to neurovascular coupling might depend on the context of the sensory stimulation. This finding is in line with the many observations of how astrocyte activity depends on the brain states.

Until now, most studies of NVC in the whisker barrel cortex have used a forced stimulation protocol. In anesthetized mice, this may be electrical (Wang, Lou et al. 2006, Lind, Brazhe et al. 2013, Mishra, Reynolds et al. 2016) or in awake stimulations, an air puff or mechanical whisker deflection (Ding, O’Donnell et al. 2013, Stobart, Ferrari et al. 2018, Tran, Peringod et al. 2018, Sharma, Gordon et al. 2020, Renden, Institoris et al. 2024). In some previous studies of evoked NVC, the duration of rhythmical deflections of the whiskers has, at times, been excessively long (Lind, Brazhe et al. 2013, Institoris, Vandal et al. 2022). We did not find self-initiated whisking that lasted longer than 10 seconds without concurrent locomotion. It implies that the results from experiments using lengthy whisker stimulations do not reproduce a natural whisking activity. In addition, this approach is also problematic because it lacks the volitional aspect of whisker-based sensing. Several intracerebral signaling mechanisms may alert the sensory cortex to an impending sensory input and could thus contribute to the NVC response. The intracerebral signaling during whisking includes synaptic input directly from the whisker motor (Kinnischtzke, Simons et al. 2014) and the prefrontal cortex (Eggermann, Kremer et al. 2014, Collins, Francis et al. 2023). Sensory responses have classically been understood in the context of enhanced local synaptic activity (Devor, Ulbert et al. 2005), but recent work has revealed that activity in adrenergic projections correlates with even the shortest whisking (Collins, Francis et al. 2023). Hence, with the intake of volitional sensory information, the subsequent need for processing might be anticipated by the barrel cortex, and both its initiation and handling depend on arousal state (Poulet and Crochet 2018). The implications are that the NVC response during whisking may change depending on whether it is for simple surveillance or coordinated exploration, as it is affected by the alertness of the animal (Shimaoka, Harris et al. 2018). Until now, only a few studies have looked at the vascular responses evoked by volitional whisking of the mice (Winder, Echagarruga et al. 2017, Tran, Peringod et al. 2018, Bojarskaite, Bjornstad et al. 2020). All but one of these focused on transitions between periods with more or less frequent whisking events instead of at the initiation of individual whisking events. These transitions reflect changes in the general behavioral states, such as from sleep to awake or quiescence to generalized activity during running. Only one previous study has, in a similar manner, correlated neuronal activity and CBF changes in the cortex of unstimulated, naturally whisking resting mice (Winder, Echagarruga et al. 2017). Our study expands on this observation by distinguishing between whisking of different durations as a measurement of the amount of sensory input. We also reveal the position of the neuronal activity and include the astrocytic contribution.

We found that astrocyte Ca2+ activity was seen to fall in two peaks: one preceding and one following the vasodilations. The first astrocytic Ca2+ peak, independent of whether the whisking was evoked, volitional during rest, or volitional prior to locomotion, reflected neuronal activity and indicated astrocytes’ reception of synaptic transmission. This activity occurred within small ROIs that we had identified through a context-dependent analysis, relying on response timing and recurrence rather than response size. The Ca2+ activity measured within these ROIs was not randomly distributed but followed a pattern of spatial reproducibility that supports that these regions are indeed biologically relevant. Because the astrocytic activity within these ROIs correlated with the neuronal activity, both with regards to distance from the vessel and response size during different kinds of whisking, we propose they are the result of activation of perisynaptic processes reacting to neurotransmitter release (Santello, Toni et al. 2019, Arizono, Inavalli et al. 2020). Their timing relative to dilation onset is essential in determining what regulatory role astrocytes could play in the NVC response. The presence of an earlier peak reflects that the astrocytic response to synaptic activity during sensory-based excitation is early enough to contribute to vasodilatory control. A relationship between synaptic activity and astrocytic Ca2+ response has been proposed by numerous studies (Bindocci, Savtchouk et al. 2017, Lind, Jessen et al. 2018, Stobart, Ferrari et al. 2018, Arizono, Inavalli et al. 2020, Armbruster, Naskar et al. 2022), but never before shown in the context of awake, volitionally whisking animals. The primary obstacle in the investigation of astrocytic contribution to NVC has been detecting the relevant activity amid all the other intracellular Ca2+ fluxes. Because we achieved this, we could investigate our hypothesis further. We did this by perturbing the astrocytic activities with a pharmacological elimination of the LC-dependent NA release (Iannitelli, Kelberman et al. 2023). This intervention did generally not eradicate the astrocytic early responses, supporting their dependence on other and possibly synaptic neurotransmitters(Cahill, Collard et al. 2023). Still, depletion of the noradrenaline levels led to attenuation of the early component of vessel-adjacent astrocyte Ca2+ activity in self-initiated whisking. Neuronal activity and PA vasodilation were not disturbed by NA depletion. Meanwhile, volitionally triggered NVC was significantly impaired, as there was no longer vasodilation of 1^st^ order capillaries, the first point of entry for local, intraparenchymal vascular inflow. Such dependence is in line with previous works showing astrocytic regulation of contractile capillaries(Biesecker, Srienc et al. 2016, Mishra, Reynolds et al. 2016, Krogsgaard, Sperling et al. 2023). It also reveals that the NVC depends on astrocytic Ca2+ when sensory input is anticipated.

We focused our investigation on key regulatory junctions in the vascular inflow tract (Grubb, Cai et al. 2020, Hartmann, Coelho-Santos et al. 2022). Here, the penetrating arteriole (PA) dives into the cortex to sustain blood flow delivery to deeper layers. Meanwhile, along its descent, 1^st^ order capillaries branch off, ensuring local and possibly layer-dependent parenchymal blood supply. We found whisking duration was a strong indicator of the size of the NVC response in the PA and 1^st^ order capillary. The correlation between whisking duration and both vessels’ dilations, as well as the activity in the surrounding neuronal structures, confirmed an obvious NVC response, even in the shortest whisking events. The spatial relationship of vessel-proximal, NVC-related neuronal and astrocytic activity has not been described before. We identified that smaller neuronal structures (<1µm^2^) closest to the penetrating arteriole (<30 µm) show the best correlation to NVC. We also found pre-emptive neuronal Ca2+ activity in volitional but not evoked whisking. A response that did not appear to influence the timing of the NVC response. Still, the early neuronal activity may be the basis of astrocytic responses that facilitate capillary dilation during volitional whisking. The response may be a layer II/III-specific phenomenon, as this layer is excited after the initial thalamocortical innervation at layer IV(Petersen 2019). Thus, the activity we detect in our imaging plane could reflect stronger cellular and vascular responses in other, deeper layers in the barrel cortex. Another unknown is the neuronal population dynamics during the different whisking events. Under the synapsin promoter, the Ca2+ indicator is expressed in all neurons, and it is not possible to distinguish between Ca2+ responses from excitatory and inhibitory neurons. Such distinctions would be relevant when measuring the degree of excitation because different neuronal populations contribute differently to NVC regulation (Howarth, Mishra et al. 2021). In addition, the groups of neurons recruited in whisking change with the context of the whisking, including locomotion(Petersen 2019). We found a strong correlation between whisking activity duration at rest and neuronal Ca2+ responses. Meanwhile, we did not see that it initially increased during locomotion. These observations are in opposition to previous reports (Ayaz, Stauble et al. 2019). This being said, our investigation focused on synaptic-size activity, disregarding the neuron cell entity, whereas many imaging-based investigations quantify cell soma participation. These differences may explain the mismatch we saw between neuronal activity and vascular responses during running. Importantly, neither was sensitive to lowered cortical NA levels. This result is supported by the recent discovery that activity in adrenergic projections does not correlate well with the onset of locomotion (Collins, Francis et al. 2023). Still, our measurements are not able to explain the size of the vascular response during locomotion, with the responses in either the neuronal or astrocytic compartment. The solution to this may be found closer to the vasculature as we did not investigate neuronal activity directly at the PA (Zhang, Ruan et al. 2024) or in the astrocytic end-feet at the sphincter.

Our results are the first to show that the earliest astrocytic Ca2+ activity can play a role in blood flow recruitment during volitional NVC. This observation suggests more dependence on astrocytic Ca2+ activities in volitional excitation over experimenter-evoked NVC, which could be meaningful in the context of the emerging understanding of the astrocyte as a regulator of changes in brain states and thus in shifts in attention (Oe, Wang et al. 2020, Reitman, Tse et al. 2023). The reduced levels of NA following the DSP-4 treatment may, therefore, impair sensory perception during whisking, which could contribute to attentional deficits to the sensory input (Cerpa, Piccin et al. 2023). An indication that this might be the case was the longer delay from whisking onset to locomotion onset in the DSP-4 treated animals (Figure 5N). Diminished vasodilation in 1^st^ order capillaries during volitional whisking both at rest and preceding locomotion (Figure 2, Figure 5O) and the resultant temporally insufficient local blood supply may also mechanistically or cooperatively hinder the sensory processing of the mouse(Devilbiss 2019). Because DSP-4 targets so broadly, other factors may also affect the lag from whisking to locomotion onset. Depletion of the global NA may suppress the higher-level decision to begin locomotion(Cerpa, Piccin et al. 2023), which might, to some degree, depend on a decreased sensory processing in Layer II/III of barrel cortex (Petersen 2019). Still, the delayed initiation of locomotion may be explained by the loss of projections from LC directly to the spinal cord or other brain regions following the indiscriminate effect of the neurotoxin(Witts, Mathews et al. 2023).

To summarize, we found that both astrocytic, neuronal, and vascular compartments reacted differently between volitional whisking, evoked whisking and locomotion. Our study indicates a novel role for astrocytes in regulating these dynamics. We also found the NVC responses were differentially affected by NA depletion between these three different types of whisker deflection. This finding has implications for understanding how patients with noradrenergic and neurovascular aberrations may respond to sensory input in different brain states and emotional contexts. It also highlights the importance of neurovascular models that encompass relevant behaviors and contexts. The perspective of this finding is that the energy support during the processing of sensory information depends on how prepared we are for the stimulation and whether we are moving or not.

## Materials and Methods

### Animals

All procedures were performed in compliance with the guidelines from the European Council’s Convention for the Protection of Vertebrate Animals Used for Experimental and Other Scientific Purposes and approved by the Danish National Ethics Committee. All experiments were performed on either WT C57bl/6j-, VGAT-ChR2-EYFP-, PV-ChR2-EYFP- or GLAST-GCaMP6-mice all of a C57bl/6j-background. GLAST-GCaMP6 mice were injected IP with 100mg/kg tamoxifen in corn oil five days in a row to induce expression of the transgene. Except for the GLAST-GCaMP6 mice, all animals were injected with either 3×200nL AAV5-gfaABC1D-lck-GCaMP6f or AAV9-syn-jGCaMP7f. Both male and female animals were used. Animals were group-housed both prior to and after chronic window implantation. Animals had access to food and water *ad libitum* and were kept on a 12/12 hour night/day cycle. Animals were at least 8 weeks old at the time of chronic window implantation, and all experiments were performed between the ages of 10 and 38 weeks.

### Chronic window implantation and viral vector injection

Chronic window implantation procedures were as in(Bindocci, Savtchouk et al. 2017). Both chronic window implantation and viral vector injection procedures were performed under aseptic conditions and isoflurane anesthesia (5% induction, 1.5% during surgery in 10% O_2_ in air mixture). Before surgery, animals were injected with dexamethasone (4.8 mg/g BW; Dexavit, Vital Pharma Nordic). The fur on top of the animal’s head was shaved off, and animals were placed in a stereotaxic frame on a heating pad to maintain 37°C body temperature. Eyes were kept moist using eye ointment (Viscotears, Novartis). The shaved skin was disinfected with chlorhexidine/alcohol (0.5%/74%; Kruuse). Carprofen (5 mg/kg BW; Norodyl, Norbrook) and buprenorphine (0.05 mg/kg BW; 19 Temgesic, Indivior) were administered subcutaneously as was a dose of 0.5mL saline distributed over several sites to keep the mouse hydrated during surgery. Lidocaine (100 µL 0.5%) was subcutaneously injected under the scalp. The scalp was removed, and the surface of the skull was cleaned. A small burr hole was drilled within the intended site for the craniotomy, and three doses (200nL) of AAV were deposited using a glass micropipette. Subsequently, a craniotomy was performed over the right somatosensory cortex (3 mm lateral, 0.5 mm posterior to bregma; Ø=3 mm). The bone flap was carefully removed, and the exposed brain was temporarily covered with a hemostatic absorbable gelatin sponge (Spongostan®, Ferrosan, Denmark) pre-wetted with ice-chilled aCSF. The cranial opening was filled with aCSF, then sealed with an autoclave sterilized round imaging coverslip (Ø=4 mm, #1.5 thickness; Laser Micromachining LTD). The rim of the coverslip was secured with a thin layer of GLUture topical tissue adhesive (Zoetis), and a lightweight stainless steel head plate (Neurotar) was positioned on the top of the skull surrounding the cranial window. The skull was coated with adhesive resin cement (RelyX Ultimate, 3M) to secure the exposed bone, including the skin incision rim, and to attach the metal plate to the head firmly. After the surgical procedure, the animals were put in their cage on a pre-warmed heating pad to wake from anesthesia. They were provided with pre-wetted food pellets for easy chow and hydration. Postoperative care consisted of subcutaneous injections of Buprenorphine 0.05mg/kg/4 times per day for up to 3 days and Carprofen 5mg/kg/1 per day for up to 5 days. The convalescent animals were carefully monitored during the 7 days of post-surgery. When fully recovered from the surgery, habituation training began. The animals were gradually habituated to the experimenter, human handling, mobile homecage (Neurotar), head fixation, and imaging. Sugar water was used as a reward during training.

### Pharmacology

DSP-4 (Tocris) was dissolved in saline, and a single dose of 75mg/kg body weight was administered at least >48 hours before imaging.

### Fluorescent dyes

All animals were injected retro-orbitally under isoflurane anesthesia 2.5% weight/volume solution of Texas-Red dextran 70kDa in saline. The injection was done at least 20 minutes before imaging to let the mouse recover fully from the anesthesia.

### Two-photon imaging

All experiments were carried out on one of two imaging systems. The first microscope was a Leica SP5, DM6000 CFS multiphoton upright laser scanning microscope equipped with a MaiTai HP Ti:Sapphire laser (Millennia Pro, Spectra-Physics) using a 20x magnification and 1.0 numerical aperture water-immersion objective. Light emitted by the tissue was split by a dichromatic filter at 560 nm into two channels and collected by two separate multi-alkali photomultiplier tubes (PMTs) after 500-550 nm and 565-605 nm band-pass filtering (Leica Microsystems). The frame size for the Leica microscope was 256 x 256 pixels, representing a physical area of 129 µm^2^. Images were collected in a single plane by bidirectional scanning with a color depth of 16 bits. The total frame time for the microscope image capture corresponded to approximately 96 ms, with a sampling rate of around 10.38 Hz.

The second microscope was a commercial laser scanning multiphoton excitation system (FluoView FVMPE-RS, Olympus) paired with a Mai Tai HP Ti:Sapphire laser (Millenia Pro, Spectra-Physics) using a 25x magnification and 1.05 numerical aperture water-immersion objective (Olympus). A dichroic mirror split emitted light with a 500-600 nm band-pass. Light reflected by the dichroic mirror was sent to a pair of gallium arsenide phosphide PMTs, corresponding to the fluorescent signal from the vasculature. Light passing through the dichroic mirror was sent to a pair of multi-alkali PMTs, corresponding to the jGCaMP7f fluorescence from the neurons. Frame size was fixed at 512 x 512 pixels, representing a physical area of 127 µm^2^. Images were collected as a single plane by bidirectional scanning using a resonant scanner, reducing the color depth to 10 bits. The total frame time was also greatly reduced to 33 ms, corresponding to a sampling rate of 30 Hz.

Images were acquired from cortical layers II–III (100–250 μm brain surface). Laser power was kept below 35 mW for all imaging to prevent phototoxicity. Laser emission was set to a wavelength of 920 nm for maximal excitation of the jGCaMP6f, jGCaMP7, and Texas Red fluorophores.

During imaging, the general behavior of the animal was recorded using a small standard video camera (Lifecam Studio, Microsoft), where the built-in IR filter had been removed and a long pass filter fitted (IR 930nm, Asahi Spectra). The LifeCam images were stored as uncompressed AVIs.

### Whisker stimulation

Air puff whisker stimulation was delivered using a PV830 Pneumatic PicoPump from WPI. The air puff was directed towards the whiskers so that there was visible deflection without hitting anything else. The pressure was adjusted to the minimum level, which still resulted in visible deflections. The stimulation consisted of 100 ms pulses delivered at 3 Hz for 2 seconds.

Enlarged whisker deflection during natural whisking was accomplished by moving a small piece of sandpaper into the trajectory of the whiskers. This was done using a custom-built motorized arm controlled via an Arduino Uno.

### Whisker tracking

An area-scan camera (GO-5100M-PGE, JAI A/S, Denmark) monitored the whisker pad and pupil, which was synchronized with the 2-photon acquisition through the same trigger circuit as the microscope. This camera was fitted with a long-pass filter to only capture infrared light (930 nm, Asahi Spectra, Japan). JAI images were stored as uncompressed 8-bit greyscale BMP images. Voluntary whisking events were detected by applying a Sobel filter to video recordings of the whiskers (Sup. Figure S1). The Sobel operator was applied to all images using Matlab’s built-in imgradient function, and each two-dimensional intensity gradient frame [i] was then subtracted from the previous frame [i-1] to determine if the image edges had changed position. The sum of the difference of all edges relative to its preceding frame was stored as the magnitude of whisker movement at that frame. The magnitudes of whisker movement were then thresholded at the median plus the standard deviation throughout the recording. Any frame above this threshold was labeled as “active whisking”. If individual events occurred within half a second of one another, the events were grouped as one single event.

### Pupil diameter detection

The JAI camera aimed at the whisker pad also allowed for accurate pupillometric measurements. A group of animals was studied separately from 2-photon imaging to ensure optimal lighting for accurate pupil detection. We quantified the diameter changes using a custom-written ImageJ script (Supplemental Figure S8). First, the pupil was cropped, and the cropped image was denoised with a median filter. The background was uniformed using the Image Calculator feature. This involved combining the pixel intensities of two images: the inverted, Gaussian blurred image and the original image. After that, edges were detected using the “Find Edges” command, and the images were binarized using local thresholding methods (Mean, Niblack, Phansalkar). To address any gaps in objects (pupils), we employed “Open” and “Filll holes” commands. Lastly, the pupil was outlined using “Analyze Particles” command. Using Hull and circle plugin, Convex Hull area, diameter of the bounding circle, and other measurements of the outlined pupils were collected.

### Movement tracking

Movement was tracked using the built-in sensor in the Neurotar Mobile Homecage system and the accompanying software (Locomotion Tracker Version 2.2.0.14, Neurotar, Finland). Movement tracking was saved as TDMS files and opened in Matlab. The listed acquisition speed was 80Hz, although a closer inspection of the timing intervals for each frame showed a hardware-level acquisition speed of approximately 77Hz. Displacement speed was thresholded at 1 cm/s to binarize locomotion.

### Data analysis of two-photon images

All final image analysis was performed using Matlab. Simple, exploratory processing was done with ImageJ, LAS AF, or Olympus software. Leica microscopy files were initially evaluated and eventually exported using Leica’s LAS AF software. The images were saved as TIFF images, and the metadata was saved in a separate XML document. Olympus microscopy files were opened, subtracted, and inspected with either ImageJ or Matlab, and all final analyses were performed in Matlab. To avoid data truncation, all image arithmetic performed on the original uint16 files, was carried out as either 32- or 64-bit floating-point operations, single and double-precision, respectively.

Motion correction: All fluorescence recordings were motion-corrected before further analysis to remedy the slight disturbances introduced by mouse movement during imaging. Although the head-fixation set-up was quite stable and even allowed for imaging during periods of significant locomotion, all experiments benefitted from movement correction (Krogsgaard, Sperling et al. 2023). We corrected for movement within and between imaging sequences to maintain the exact field of view. Earlier movement correction was cross-correlation based, but we found normalized 2D cross-correlation corrected even finer movements.

Vasodilation: Vasodilations were calculated as dA/A. Penetrating arterioles and 1^st^ order capillaries were manually selected and thresholded. Fluorescence videos of blood vessels were spatially smoothed using a 3 by 3 pixel median filter, preserving sharp edges while denoising with a Gaussian filter. Masks for vessels of interest were drawn by hand. All frames were thresholded using the triangle thresholding method. The binary images were cleaned up using morphological opening and closing, and the foreground area was calculated. The median of the whole trace was used as a baseline level to normalize dilation traces. The area was calculated and normalized by the median of the area trace for an entire recording. Using the triangle method, we thresholded and binarized the imaging sequence based on its intensity histogram. Morphological clean-up was performed, and the remaining binary area was taken to be the vessel area. This process was repeated for first-order capillaries as needed, with the only difference being the capillary ROI was manually outlined and applied as a polygon. Finally, the remaining binary areas were then used to determine the frame-by-frame change in area of the respective vessels and informed the perceived vasodilation and vasoconstriction. The median of the whole trace was used as a baseline during normalization.

### Ca2+ ROI detection

#### Astrocytic dynamic ROIs

Ca2+ microdomain detection: We developed a method that looked at changes in fluorescence and not the fluorescence strength. The method consisted of first thresholding, then cleanup of active voxels (two spatial dimensions and one temporal) – then clustering of detected active voxels – and then analysis of these clusters (Sup. Figure S3). The detection scheme outlined below is not too dissimilar to AQuA (Wang, DelRosso et al. 2019), although there are key differences. Thresholding was based on two parameters - the baseline (µ) estimated with the mode and the baseline variation (σ). A mode of the raw trace underestimates the baseline level as many pixels would often show no signal and, therefore, lie at the minimal fluorescence intensity value possible. Instead, we used the value around which the signal fluctuates at rest. This value was found on smoothed data since, in this step, we are only interested in the level around which the signal fluctuates and not the fluctuations themselves. Hence, signals from each pixel were lowpass-filtered using a 5^th^ order Butterworth filter with a cutoff frequency of 0.26 Hz (5% of the Nyquist rate)(Sup. Figure S3B). Then, the data was binned to width 50 counts, and the fluorescence signals were in the order of 10000s. This binning sacrifices some resolution to circumvent the issue of the minimal value appearing several times. By binning the signal, the value around which the signal fluctuates should be better represented as the single minimal value around which the signal does not fluctuate. After filtering and binning, the mode was found to estimate the baseline level. An estimate for the baseline variation around the baseline level was found by high-passed filtering of the signals again using a Butterworth filter with the same cutoff frequency as the lowpass filter. By doing this, we discard the larger changes stemming from actual Ca2+ activity within the cell and just get the high frequency of noise through which the signals must stand out to be detected. As a first rough estimate of the baseline variation, we take the standard deviation of this high-pass filtered trace. Finding the variance using the standard deviation of this signal will be an overestimate, as the fluorescence noise is partly due to Poisson processes in the fluorophores and in the detectors. For the Poisson distribution, the variance is equal to the mean (µ = σ2), so with more signal strength, we get larger variation as well. Thus, we used a variance stabilization strategy based on iterations. To find the baseline, we threshold the signal with our current estimates (the baseline level is found as described above, and the baseline variation is represented by the standard deviation of the high pass filtered signal).

Anything below a threshold of µi + 2 · σi the i’th pixel) is considered to belong to the true baseline. We then used this new trace, where we discarded a large portion of the Ca2+ signals, to better estimate the baseline level and variation (Sup. Figure S3D). We then used this estimation of baseline level and variation to threshold each pixel signal (Sup. Figure S3E). Next was a clean-up of the detected active voxels. Even after a threshold at two sigma, there will still be some false positives; thus, lone voxels without any active neighbors were disregarded, and voxels that were close together were grouped. Since the data is a binary 3D set (spatiotemporal, Sup. Figure S3F), this can be accomplished by using simple morphological operations known as opening and closing, commonly used for noise removal in computer vision (Sup. Figure 3G).

The next step clusters the active voxels into domains that are active together. Clustering was done in 2D, but to avoid clustering events that might partially or wholly be within the same spatial region but do not overlap in time, the binary 3D array was first projected into 2D. The overlapping clusters were thus projected onto the same or connected pixels and consequently clustered together. This may result in an underestimation of the amplitude of spatially small signals that happen to lie within a larger cluster as the average fluorescence of the whole area is calculated for each frame. If only part of the cluster is active in some frames, this will lead to underestimation of that signal.

This metric yields the total area of the FOV, which was active at some point during the recording, the total integrated fluorescence of all signals within the recording, and the number of spatial regions where the activity took place. ΔF/F-traces from these clusters can be further worked upon (Sup. Figure S3I). The accuracy of this detection algorithm was tested on synthetic data produced at various levels of contrast and signal-to-noise ratio (SNR).

#### Astrocytic context-dependent ROIs

To identify regions of astrocyte activity in relation to NVC processes, we first detected dilation peaks in the PA area trace (Sup. Figure S5). Dilation events preceded by whisking but did not overlap with locomotion or air-puff stimulation were selected. Only periods with at least 2 seconds of baseline prior to the dilation peak were included. The average fluorescence levels in a 1 second period from 2-1 second before the dilation peak was averaged. From this, a mask was created by applying a 2D Gaussian filter (imgaussfilt.mat) to denoise and threshold the ROIs. Using the bwareafilt.mat function areas smaller than 1.5µm^2^ pixels were excluded. The filtered image was binarized to create a mask. In this image, we identified regions with maximal activity in the astrocytic Ca2+ levels one second before the dilation peak. These ROIs were then grouped depending on the distance from the PA. The innermost 15µm ring from the center of the PA was excluded to eliminate any disturbances from the changes in vessel diameter. That also meant that end-feet structures were not investigated in this study. The data was then grouped into four zones counting from the outer rim of the innermost circle: <15µm from PA, 15-30µm from PA, 30-45µm and 45-60µm from PA. The average activity in these ROIs was then measured for each image in the acquisition in the entire imaging session.

#### Neuronal ROI placement

To find the regions most active during whisking, we created average images of the periods surrounding whisking. We began with sorting whisking events and creating ΔF/F Ca2+ images (Supplemental Figure S7A). First, all whisking events overlapping with movement and within the first 0.2 seconds of recording were discarded. Then, based on our astrocytic data, all whisking events with durations shorter than one second and longer than three seconds were discarded to isolate hemodynamically relevant 1-3 second duration whisking events. ΔF/F baseline was extracted from the mean per pixel value of Ca2+ images. All images in the sequence were then subtracted and divided by this baseline, converting all images in the sequence into respective ΔF/F values (Supplemental Figure S7B). For the remaining one- to three-second-long whisking events, the Ca2+ images corresponding to seven seconds before and after each whisking onset were summed together to form 14-second-long cropped average Ca2+ images, which were centered on the onset of the whisking episode (Supplemental Figure S7B).

Having created whisking-aligned composite images, we defined a 500 millisecond (ms) window around whisking, 100 ms before and 400 ms after whisking onset. We normalized our composite ΔF/F images on their per-pixel distributions. We used the periods prior to whisking onset with less than 50% whisking activity as a baseline and subtracted the per-pixel baseline from all images. We then calculated the standard deviation of each pixel over the entire 14-second period and divided each pixel by its standard deviation. From these normalized images, we identified the most active regions in the half-second window (Supplemental Figure S7C).

We selected the 120 most active regions, as indicated by their respective normalized values. Next, these regions were automatically grouped into three categories according to size and shape: soma, dendrite, and synapse (Supplemental Figure S7C). We noticed small active areas without clear compartmental identities and, depending on size, labeled them as dendritic (>1µm^2^) or synaptic(<1µm^2^). These automatic morphological groupings were then manually checked and recategorized as necessary by comparing them to manually outlined soma maps and field-of-view intensity projections (Supplemental Figure S7A). We extracted traces for each imaging sequence from each ROI, three morphological fields, and two fields of view: soma, dendrite, synapse, neuropil, and field of view. This entire process was repeated for each imaging session separately due to slight variations in our z-plane between imaging days.

### Statistics

To compare the anticipatory and reactive responses of our stimuli, sandpaper, and air puff, to their respective controls, we classified whisking activity into four different categories: volitional whisking at rest with and without sandpaper, air puff evoked whisking events and whisking events that overlapped with locomotion events of >2 seconds duration. The analysis did not include whisking that overlapped with locomotions shorter than 2 seconds. For comparison between volitional whisking events during rest, with or without sandpaper, only event lasting between 1 to 3 seconds was included. When comparing to the 2-second duration air-puff, only volitional whisking events lasting 1.5 to 2.5 seconds were included. When comparing volitional whisking events during rest to those during locomotion, only events >2 seconds were included.

Responses to separate whisking events were averaged according to the FOV from which the measurements were made. The average response to all whisking events in each FOV was used for statistical testing, as they were considered the relevant entity. Namely, this specific PA, 1^st^ order capillary, astrocytic or neuronal structure. The number of spontaneous whisking events of different duration would differ between FOVs. For their number per FOV in astrocytic and neuronal mice before and after DSP-4 administration, see supplemental figure S15. N=20 FOV from 16 mice was included in the investigation of the blood vessel and astrocyte Ca2+ activity investigation. Most of these mice were also included in another study(Krogsgaard, Sperling et al. 2023). N=19 FOVs from 7 mice were included in the study of neuronal Ca2+ activity.

The baseline was calculated per event as each whisking event was individually normalized to the activity directly preceding it. The trace one to eight seconds before each whisking onset was taken as a baseline for each whisking event. Periods within two seconds of movement were omitted. For air puff trials, we found the limited number of trials made the traces and respective baselines more susceptible to excessive animal movement. To address this, we shortened the baseline period to one to four seconds before stimulus onset. This reduced trace variance before and after stimulus onset.

Statistical tests were applied depending on the type of data. If normality could be expected, it was tested visually with a qq-plot, and data were log-transformed when appropriate. Wilcoxon signed-rank test for paired and Mann-Whitney for unpaired data was used when comparing time-locked or counted data as these could not be expected to show normal distribution. When data was normally distributed the following tests were used: Two-tailed paired T-test; Welch’s t-test if there was a large difference in N size between the two tested groups as was the case when comparing air puff against 2-second whisking events; One-sample t-tests were used to compare traces to the baseline; Mixed-effect linear model test was used instead of paired t-tests when comparing stimuli to their controls: sandpaper against free whisking and Drug effect on volitional whisking. The Mixed model was used because some FOVs came from the same mouse and thus could not be considered independent. When investigating multiple comparisons, Mixed effects models were also used to test significance, followed by two-tailed paired or unpaired Student’s t-test as appropriate and finally with Holm-Sidak correction for multiple comparisons. Significance was set at a level of 0.05. Single stars indicate p-values < 0.05, two stars < 0.01, and three stars < 0.001.

## Acknowledgment

We would like to acknowledge our animal technician, Micael Lønstrup, for his help with animal experiments. Thanks to Dr. Kirsten Thomsen for statistical advice and guidance. Professor Martin Lauritzen and associate professor Krzysztof Kucharz for good discussions. This study was supported by Danmarks Frie Forskningsfond “1030-00374B”, “Læge Sofus Carl Emil Friis og Hustru Olga Doris Friis’ Legat”, “Dagmar Marshalls Fond,” and “Hørslev fonden”.

## Author’s contribution

B.L. and A.K. designed the study. A.K., J.S., G.K., L.S., and B.L. performed the experiments. A.K., J.S., G.K., and B.L. analyzed the data. B.L., A.K., and J.S. wrote the paper with input from all authors.

## Data availability

The data supporting these findings are available from the corresponding author upon reasonable request.

## Code availability

Code is available from the corresponding author upon reasonable request.

## Supplemental information

**Supplemental Figure S1:**
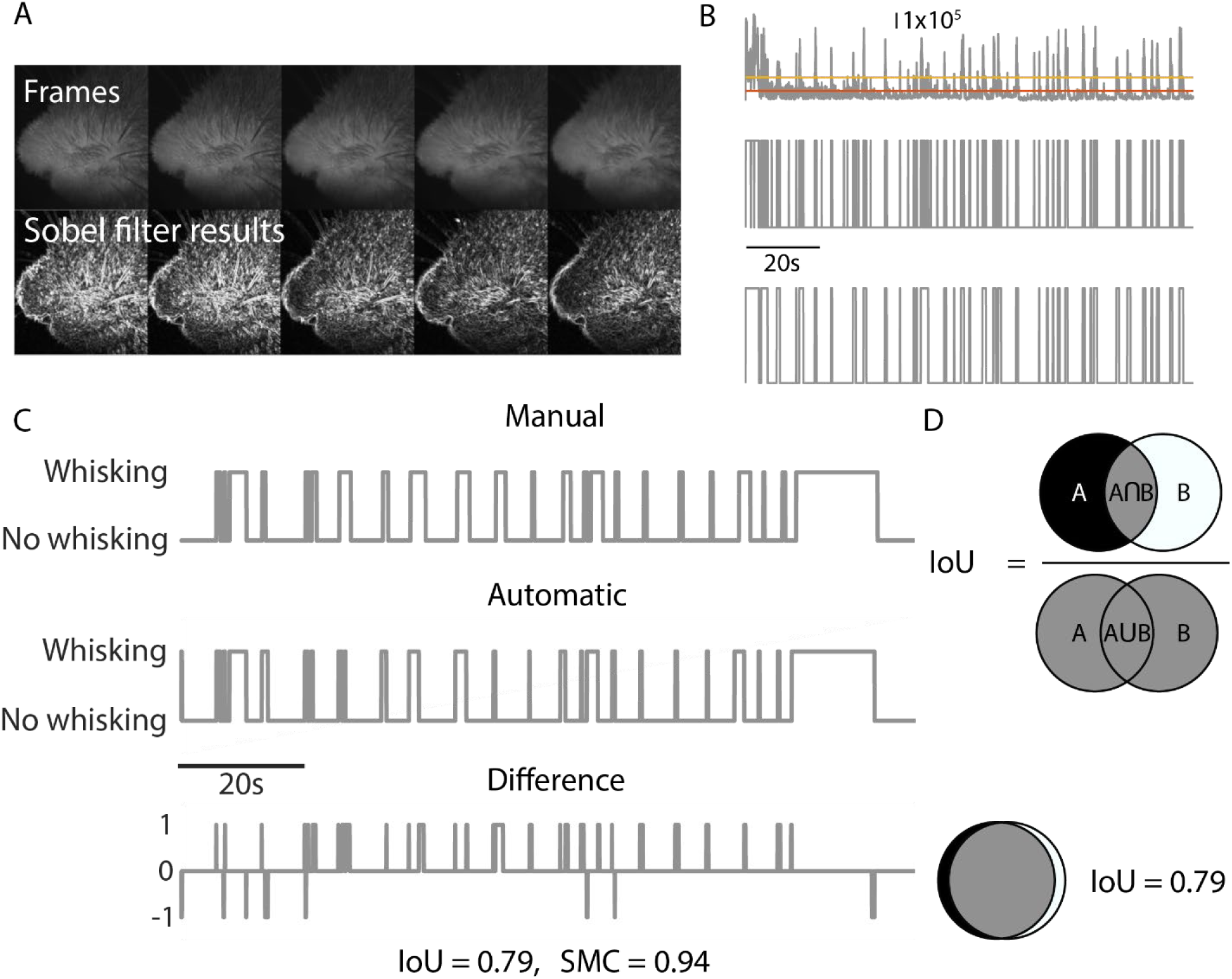
How whisking activity was evaluated from video data. **A)** A sobel filter is applied to all frames. Frame i-1 is then subtracted from frame i, yielding only differences frame by frame. The absolute value of each pixel value is taken such that both negative and positive changes in pixel value count towards a change from the preceding frame. The region containing the whiskers is cropped and sum of the cropped area is calculated. This sum represents the *amplitude* of the frame-to-frame difference, which represents a movement of the whiskers. **B)** <u>Top</u>: The sum of frame-to-frame difference as a function of time (*grey*). The median of the traces is shown as a *red line*. The *yellow line* shows the median + standard deviation. <u>Middle</u>: The raw difference traces shown at the top are thresholded by the median + standard deviation, yielding a binary trace. <u>Bottom</u>: The binary trace shown in the middle is cleaned up using morphological operations. **C)** Comparison of manual and automatic detection of whisking events. The bottom trace shows the difference between the two detected binary event traces. There is good agreement between the two traces, and disagreements are mainly about the exact start time or end time of the event but rarely about failure to detect an event using either method. Using the Intersection over Union (IoU) measure (also known as the Jaccard index) to measure the agreement between the manual and automatic detection yields a value of 0.79 (1 being identical results). Using the Simple Matching Coefficient (SMC) yields 0.94 (again, 1 being identical results) **D)** A cartoon showing the principle of the IoU measure and an illustration of what an IoU of 0.79 looks like.

**Supplemental Figure S2:**
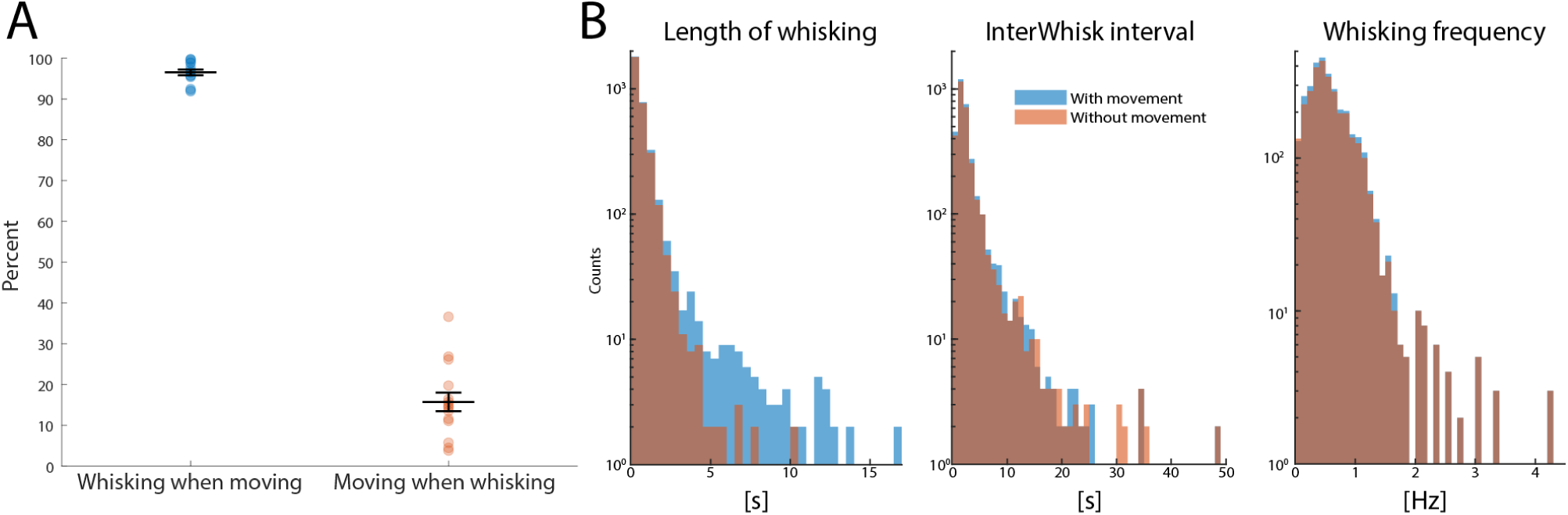
The relationship between whisking and moving. **A)** Moving is always associated with whisking but not the other way around. About 95% of the time, when the animal is running, it is also whisking, but less than 20% of the time spent whisking is also spent running. **B)** Histograms showing the distributions of whisking events, including whisking during locomotion (*blue*) and excluding locomotion-associated whisking (*red*). Whisking events longer than 5 seconds were very rare in the absence of running. The time interval between two consecutive whisking events was usually quite short, with the mode being 2 seconds. Taking the reciprocal of the interwhisk interval yields a whisking *frequency* with a mode of 0.5Hz.

**Supplemental Figure S3:**
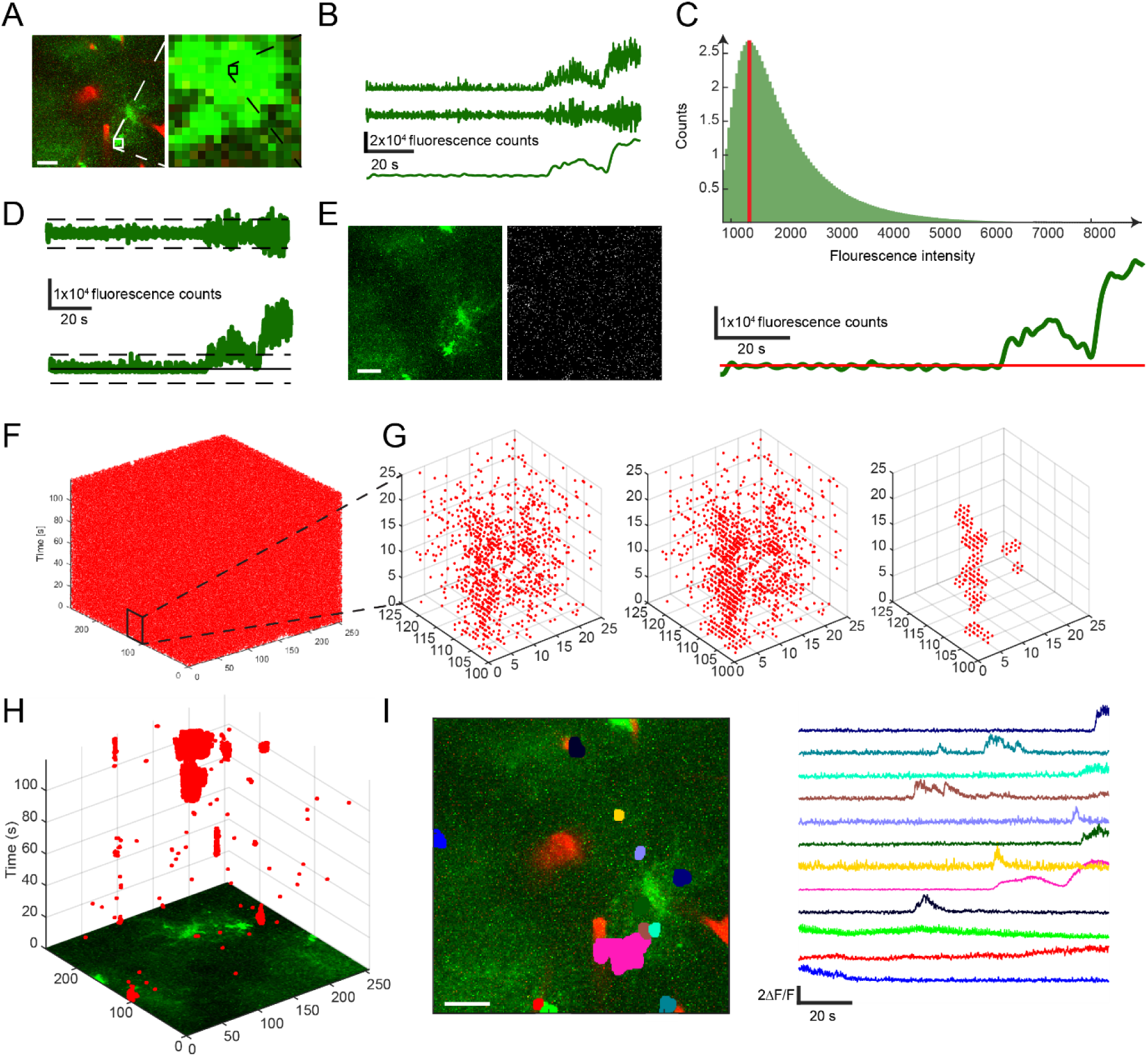
Event detection algorithm. A) Example frame from a two-photon recording. Scalebar 20µm. Each pixel is analyzed for changes in signal intensity. The detected active voxels are later combined into domains and analyzed together. B) <u>Top:</u> raw signal from the pixel highlighted in A). <u>Middle:</u> Signal after high pass filtering. <u>Bottom:</u> Signal after low pass filtering. C) <u>Top:</u> Histogram of signal values of low pass filtered signal. The mode is highlighted in *red*. <u>Bottom:</u> Mode plotted on top of low pass filtered signal. The mode is used as an estimate of the baseline level. D) <u>Top:</u> A high-pass filtered signal is used to estimate baseline variation. Standard deviation. <u>Bottom:</u> Baseline level and baseline variation estimates plotted on top of raw signal. E) All frames (example frame is shown on the left) are thresholded using estimates of baseline level and variation Scalebar 20µm., resulting in binary images (right). F) Each recording then becomes a binary 3D (xyt) array. The thresholding will inevitably result in some false positives due to noise. G) <u>Left:</u> A zoom-in on a small subvolume of the whole recording. <u>Middle</u>: Same subvolume after clean-up using morphological closing. <u>Right:</u> Same subvolume after clean-up using first morphological closing and then morphological opening. H) Resulting in a binary 3D array showing detected events. Active voxels are clustered using connected components into active domains. I) <u>Left</u>: Active astrocytic domains within a FOV were detected. Scalebar 20µm. <u>Right</u>: corresponding normalized fluorescence traces from the detected domains.

**Supplemental Figure S4:**
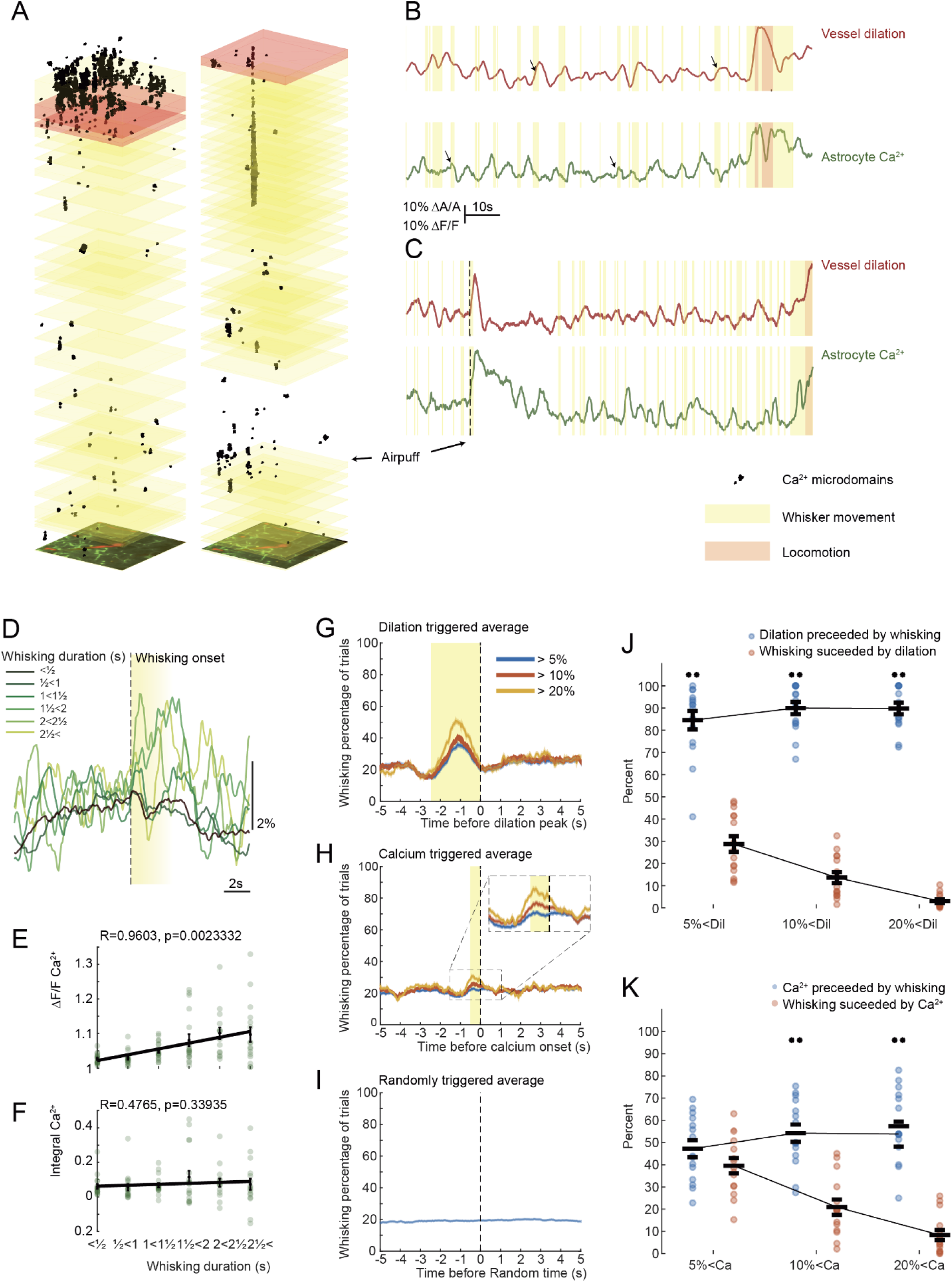
Measured with dynamic ROIS, astrocytic calcium activity was similarly but less robustly associated with whisking. The connection between natural whisking and arteriolar dilations, as well as astrocytic calcium elevations, was better than random. **A)** 3D (xyt) visualizations of astrocytic calcium events. *Yellow*-shaded areas are whisking events, and red-shaded areas are locomotion events. Large increases in calcium events are seen in response to locomotion. **B)** Vessel area trace (*red*) and astrocytic calcium activity (*green*). Arrows indicate some responses that correlate with whisking events. Traces correspond to the recording, which is 3D-visualised on the left in **B)**. **C)** Vessel area trace (*red*) and astrocytic calcium activity (*green*). Arrows indicate response to an air puff stimulation of the whiskers. Traces correspond to the recording, which is 3D-visualised on the right in **B)**. **D)** Average astrocytic calcium traces from dynamic ROIs for volitional whisking events of increasing duration. N= 20 locations in 16 mice. **E)** The amplitude of astrocytic calcium responses as a function of whisking event duration. Pearson correlation R = 0.96, p = 0.0023. N= 20 locations in 16 mice. **F)** AUC of astrocytic calcium responses as a function of whisking event duration. Pearson correlation R = 0.48, p = 0.34. N= 20 locations in 16 mice. **G)** Dilation triggered average of whisking activity for vasodilatations above 5%, above 10%, and above 20%. A clear peak is seen in whisking activity within the 2.5 half seconds before the peak of the vasodilation. **H)** Calcium-triggered average of whisking activity for calcium increases above 5%, above 10%, and above 20%. An increased whisking activity about half a second before the calcium peak for the cases of calcium increases above 10% and above 20%. **I)** The randomly triggered average of whisking activity was calculated in the same manner as the dilation-triggered and calcium-triggered average, but random timepoints were used instead of calcium or vasodilation peak times. The flat curve reflects a lack of correlation to the random timepoints. **J)** *Blue* dots represent how many vasodilations are preceded by whisking activity within the 2.5 seconds before the peak. This was tested against how many random time points were associated with whisking activity in the 2.5 seconds before the time point. In all cases (above 5%, above 10%, above 20%) significance was found (**p<0.01). *Red* dots represent how many whisking events are followed by vasodilations within the 2.5 seconds following the whisking event onset. A relatively large portion of whisking events did not evoke vasodilations. If discarding the very shortest whisking events, which represent the largest group of whisking events, the fraction is approximately the same, showing that not all whisking events trigger vasodilations larger than 5%. **K)** Same as **D)** but for the case of astrocytic calcium responses. Significantly better than random behavior was only found when considering Ca2+ responses larger than 10% and larger than 20% (**p<0.01).

**Supplemental Figure S5:**
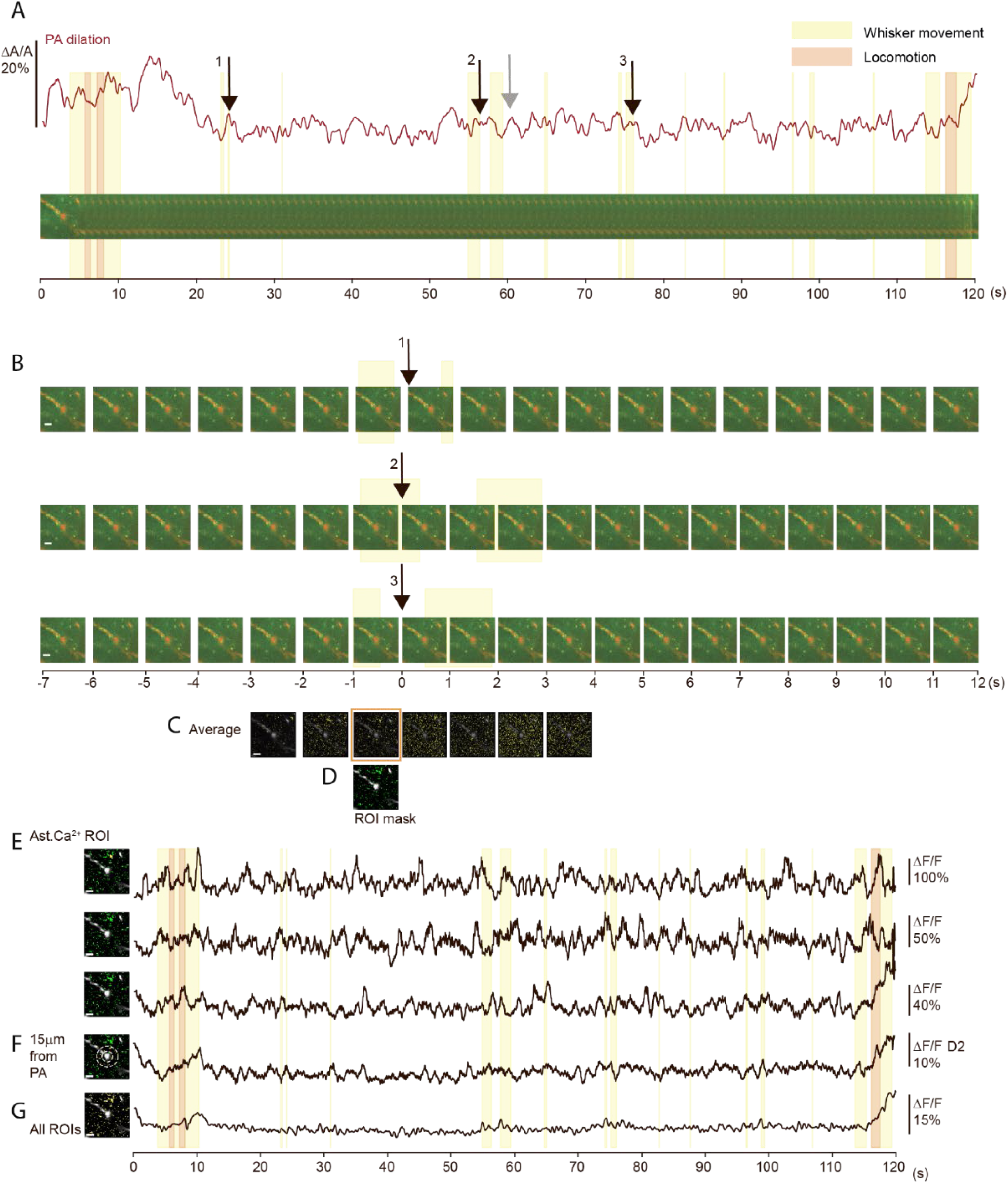
Identification of context-dependent regions of interest in astrocytic Ca2+ data. **A)** Above: PA dilation trace from a 2 min imaging session. *Black* arrows point to three dilation events preceded by whisking. *Grey* arrow points to a similar dilation event but with an insufficient baseline period to be included in the procedure of detecting context-dependent ROIs. <u>Bottom</u>: Imaging sequence with *Red*: Texas red dextran in vessel and *Green*: Glast-GCaMP6, averaged for every 1 s for illustration. Periods of whisking are shown in *yellow* and locomotion periods are in *red*. **B)** Three shorter sections of the image session in A) aligned to the peak of dilation 1, 2 and 3. **C)** Averaged images of the activity aligned to dilation peak in B) 3 seconds before and four seconds after the dilation peak. **D)** Mask created from maximal activity in the astrocytic Ca2+ levels one second before dilation peak. **E)** Traces of average Ca2+ activity from three individual ROIs during the entire imaging session. <u>Left:</u> position of the ROI. <u>Right:</u> average timeseries trace. **F)** Traces of average Ca2+ activity from all ROIs within 15µm of the center of the PA during the entire imaging session. <u>Left:</u> two circles demark the region from which the ROIs are included. <u>Right:</u> averaged timeseries trace. **G)** Traces of average Ca2+ activity from all ROIs during the entire imaging session. Scalebars 20µm.

**Supplemental Figure S6:**
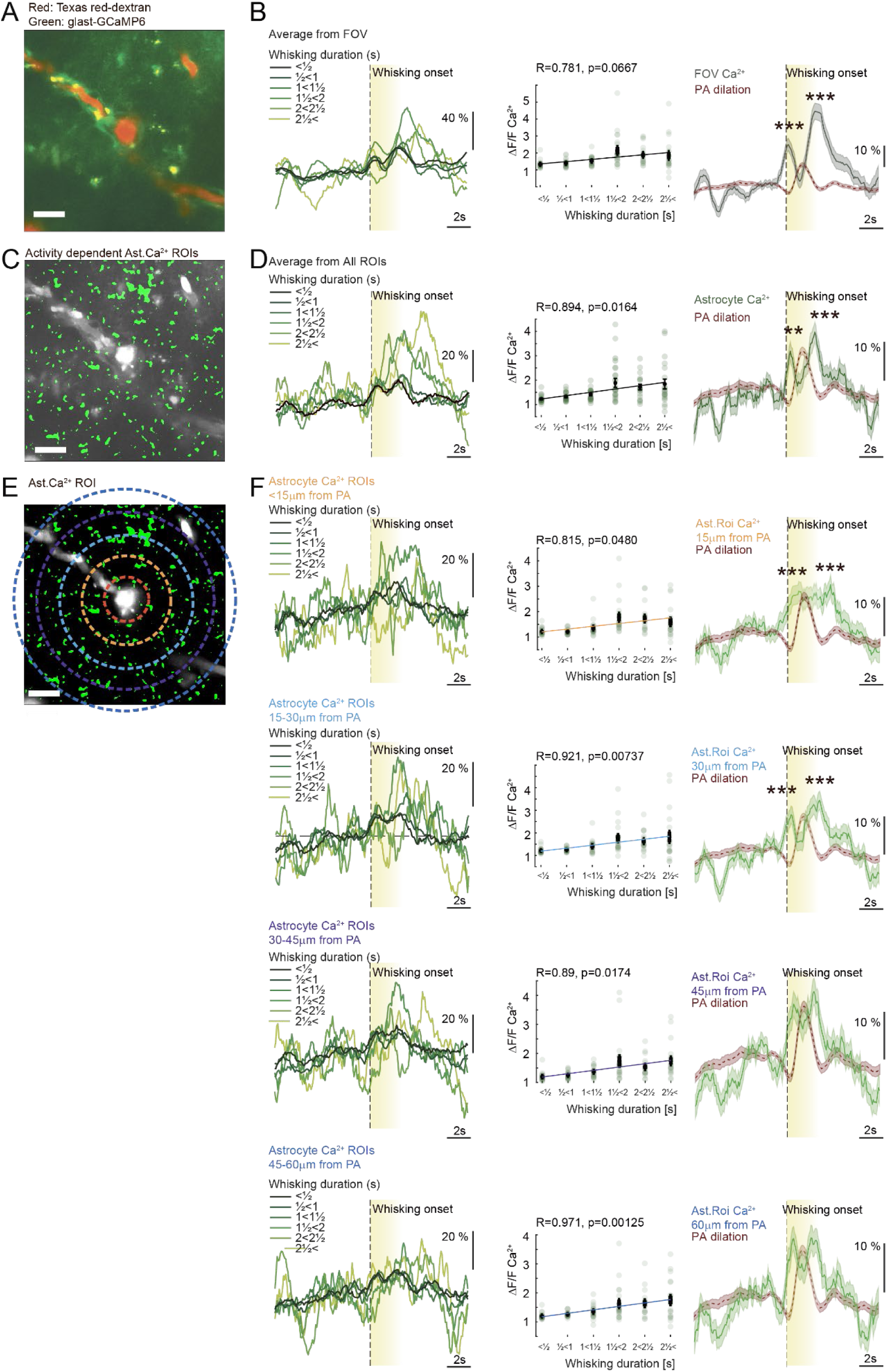
Whisking event dependent astrocytic ROI, were grouped depending on distance from PA. **A)** Average timeseries from 2-photon microscopy of vascular inflow tract with Texas red-dextran in the vessel lumen and astrocytic Ca2+ indicator GCaMP6f expressed under the GLAST promotor. **B)** <u>Left:</u> Astrocytic Ca2+ averaged in the FOV during all volitional whisking events of increasing duration aligned to whisking onset. <u>Middle</u>: Amplitude of astrocytic calcium responses as a function of whisking event duration. Pearson correlation R = 0.78, p = 0.067. <u>Right:</u> Average astrocytic Ca2+ activity in FOV during all volitional whisking events lasting >1s shows the two peaks in the astrocytic Ca2+ responses. Levels significantly above baseline both prior to and after PA dilation (*red*). Outline of 1 s.e.m. T-test vs baseline: Early Ca2+ max: p=6.5*10-124 AUC: p=0.00026, Late Ca2+ max: p=5.7*10-126 AUC p=6.1*10-13, N=989, 20 FOV 16 mice. **C)** Image from (A) overlayed the context-dependent astrocytic ROI selection. **D)** <u>Left:</u> Astrocytic Ca2+ averaged in all ROIs in C) during all volitional whisking events of increasing duration aligned to whisking onset. <u>Middle</u>: Amplitude of astrocytic calcium responses as a function of whisking event duration. Pearson correlation R = 0.89, p = 0.016. <u>Right:</u> Average astrocytic Ca2+ activity in all ROIs during all volitional whisking events lasting >1s (*Green*) shows levels significantly above baseline both prior to and after PA dilation (*red*). Outline of 1 s.e.m. Test vs baseline: Early Ca2+ max: p= 8.7*10-153 AUC: p=0.0033, Late Ca2+ max: p= 3.4*10-171 AUC p=0.00092, N=989, 20 FOV 16 mice. **E)** Image with ROIs from (C) overlayed with the 15 µm zones dependent on distance from the center of the PA used to group the ROIs. **F)** <u>Left:</u> Astrocytic Ca2+ averaged in all ROIs in the different zones in E) during all volitional whisking events of increasing duration aligned to whisking onset. <u>Middle</u>: Amplitude of astrocytic calcium responses as a function of whisking event duration. <u>Right:</u> Average astrocytic Ca2+ activity in all ROIs during all volitional whisking events lasting >1s (*Green*). T-test vs baseline **p<0.01. ***p<0.001.* show levels significantly above baseline both prior to and after PA dilation (*red*). Outline of 1 s.e.m., N=989, 20 FOV 16 mice. Scalebars 20µm.

**Supplemental Figure S7.**
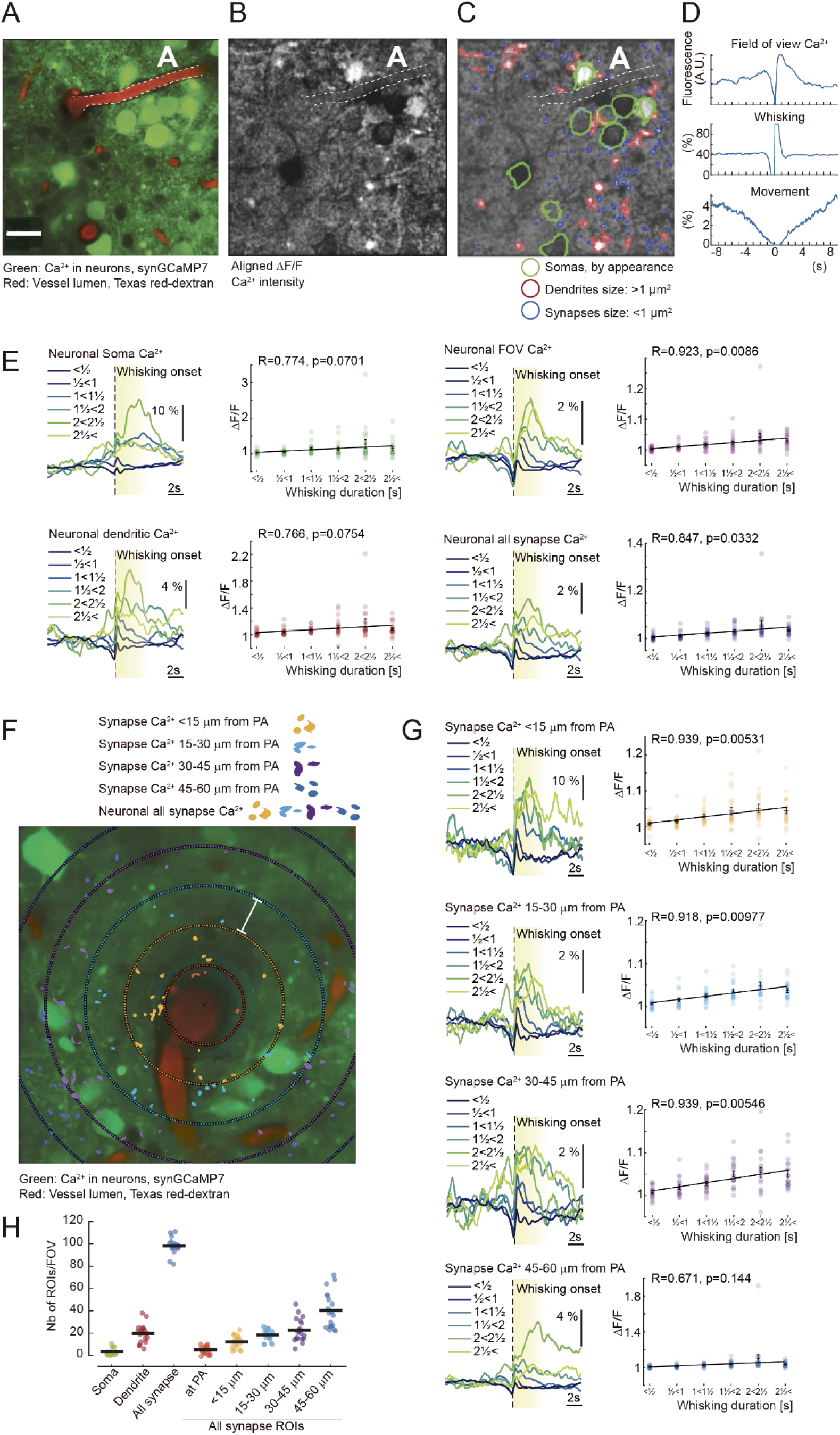
Detection of spatially specific neuronal activity using context-dependent ROI definition method. **A)** Average timeseries from 2-photon microscopy of vascular inflow tract with Texas red-dextran in the vessel lumen and neuronal Ca2+ indicator GCaMP7 expressed under the synapsin promotor. The white A and the *dashed line* marks the artery to compare with B) and C). Scalebar 20µm. **B)** Same data set as in A). Now, images are averaged centered around whisking onset (from 100 ms before whisking onset to 400 ms after whisking onset). Pixel values are given in units of ΔF/F. Based on this image, ROIs are selected. **C)** The averaged, temporally centered ΔF/F image from B) with categorized ROIs. *Green* outlines pre-traced soma boundaries. *Red* outlines dendrite-labelled structures (area >1 µm^2^). *Blue* outlines synapse-labelled structures (area <1 µm^2^). **D)** <u>Top:</u> Average field of view calcium activity within an 18 second time window. Framerate is 30 Hz. <u>Middle</u>: Average whisking activity within this same time frame. Note 0% whisking activity within the 0.5 second exclusion period before whisking onset and 100% whisking from frame 270 (whisking onset). <u>Bottom</u>: Average movement trace activity. Note 0% movement during whisking activity. **E)** Neuronal Ca2+ activity averaged within different areas: Neuronal soma, FOV, dendritic ROIs and all synaptic ROIs. <u>Left:</u> during all volitional whisking events of increasing duration aligned to whisking onset. <u>Right:</u> Amplitude of neuronal calcium responses as a function of whisking event duration and their Pearson correlation. **F)** Categorization of synaptic ROIs dependent on their distance from the center of the PA overlayed an average timeseries from 2-photon microscopy of vascular inflow tract with Texas red-dextran in the vessel lumen and neuronal Ca2+ indicator GCaMP7 expressed under the synapsin promotor. Scalebar 15µm. **G)** Neuronal Ca2+ activity averaged within different zones from the vessels with 15 µm steps. <u>Left:</u> during all volitional whisking events of increasing duration aligned to whisking onset. <u>Right:</u> Amplitude of neuronal calcium responses as a function of whisking event duration and their Pearson correlation. **H)** Number of ROIs detected in each FOV, grouped based on size in um^2^ and with regards to the synaptic-like ROIs also on distance from the PA. All data from 19 FOV in 7 mice.

**Supplemental, figure S8.**
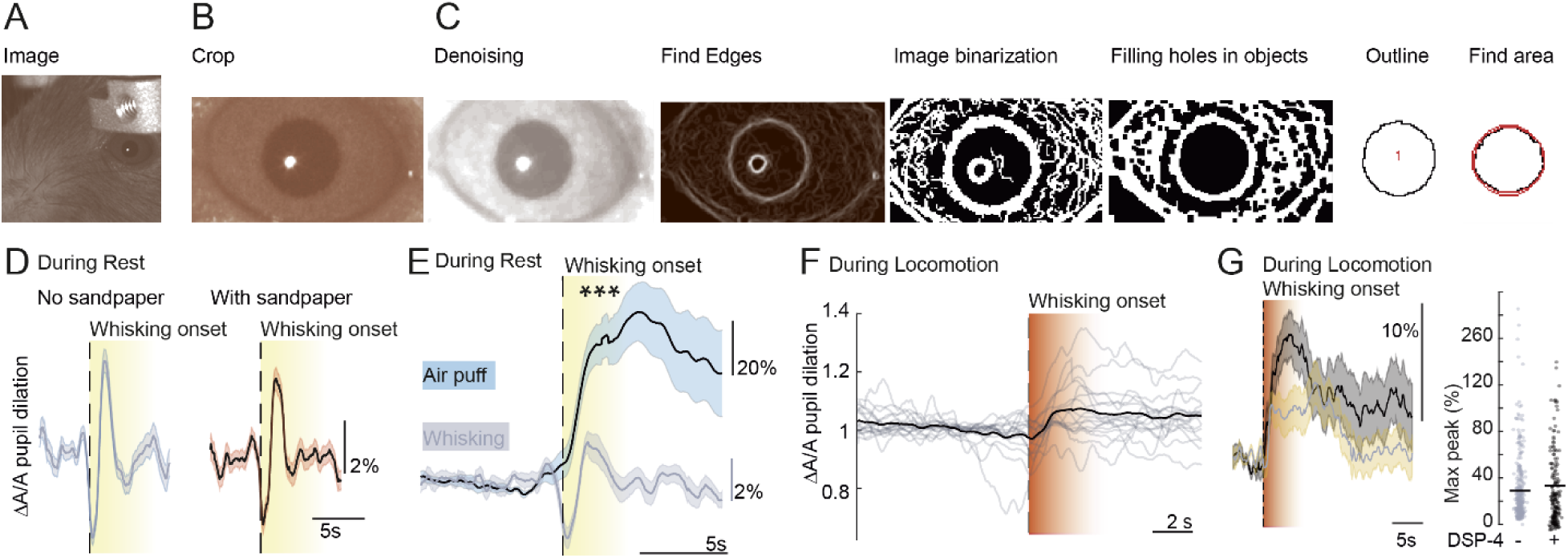
Pupil dynamics detection. **A)** Infra-red image of the head-fixed mouse. **B)** Crop from A) showing only the eye. **C)** The image of the eye was denoised with a median filter. The background was uniformed using the Image calculator feature, combining pixel intensities of an inverted Gaussian blurred image with that of the unchanged image. Edges were detected using “Find Edges” command. The image was binarized using local thresholding methods (Mean, Niblack, Phansalkar). In order to remove any gaps in the object (pupil), “Open” and “Fill holes” commands were employed. Lastly, the pupil was outlined using “Analyze Particles” command. Using Hull and circle plugin, Convex Hull area, a diameter of the bounding circle and other measurements of the outlined pupils were collected. **D)** Pupil diameter during all volitional whisking events 1½<3s long, aligned to whisking onset without (left, *grey*) or with (right, *orange*) sandpaper had no effect on the average pupil fluctuations. **E)** Average pupil dilations during volitional whisking events lasting between 1½-2½ seconds (*grey, solid line*) were significantly smaller than during 2s air-puff evoked whisking (*dashed line, black*, 1 s.e.m. outline in *light blue*). Paired t-test, p=0.000128, N=6 mice. **F)** Pupil activity aligned to whisking during locomotion. Individual responses are in *grey* and average (*black trace*). **G)** The effect of DSP-4 on pupil diameter during locomotion was not significant, T-test, p=0.12 N=6 mice.

**Supplemental Figure S9:**
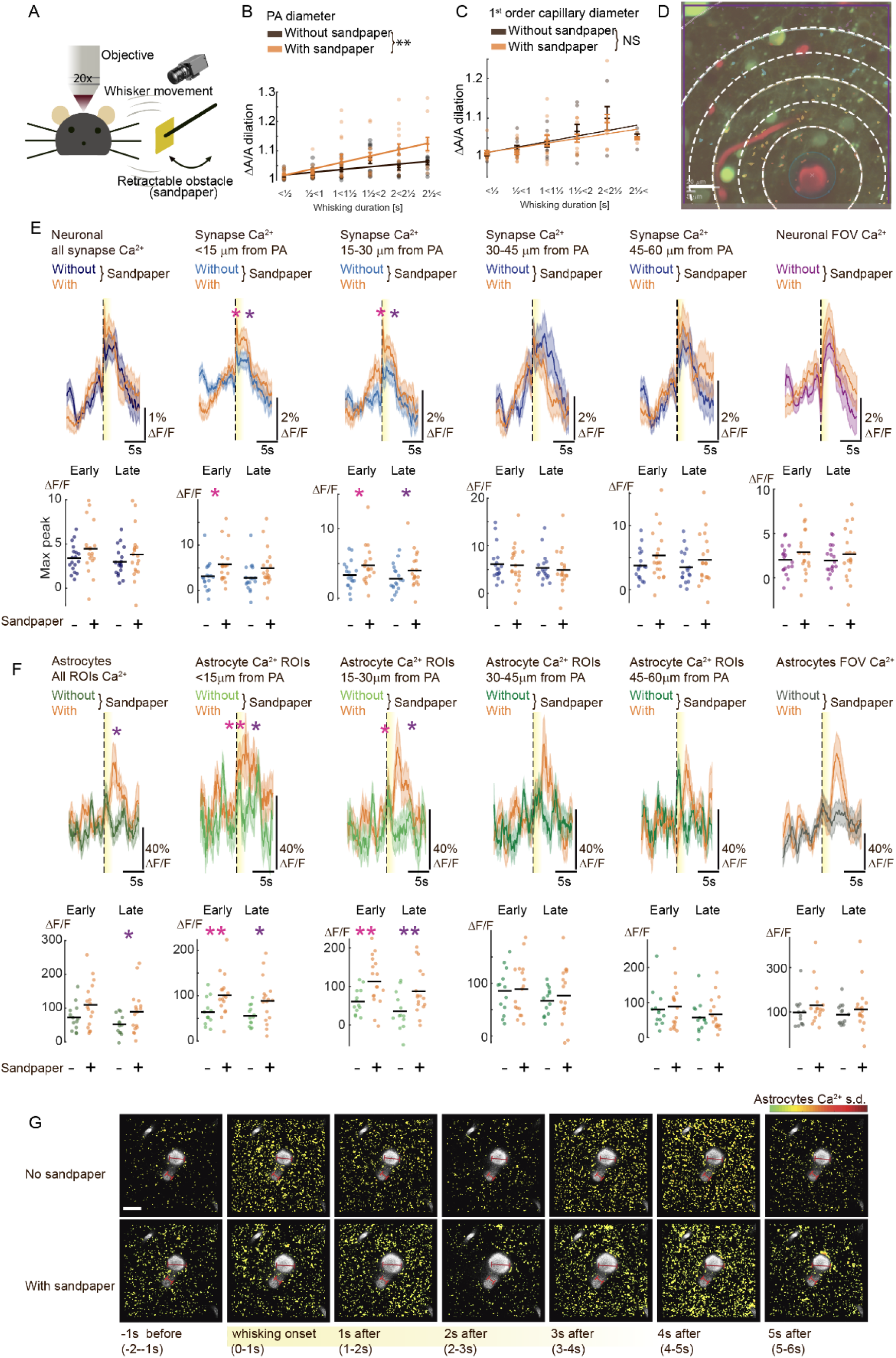
Neuronal and astrocytic Ca2+ responses without and with sandpaper sorted depending on the distance from PA. **A)** Cartoon of the setup showing the principle of the retractable obstacle. An infrared camera records whisker movements and a piece of sandpaper is connected to a motorized arm to make it retractable. **B)** PA dilation amplitude as a function of whisking duration showed significant enlargements comparing whisking without (*black*) and with sandpaper (*orange*). N= 20 locations in 16 mice ANCOVA, **p<0.01. **C)** 1^st^ order capillary dilation amplitude as a function of whisking duration showed no significant effect of the sandpaper comparing whisking without (*black*) and with sandpaper (*orange*). N= 10 FOVs in 7 mice. ANCOVA, p>0.05. **D)** Illustration of the 15µm zones sorting ROIs dependent on their distance from the center of the PA. Overlayed an average timeseries from 2-photon microscopy of vascular inflow tract with Texas red-dextran in the vessel lumen and neuronal Ca2+ indicator GCaMP7 expressed under the synapsin promotor. **E)** <u>Top</u>: Average neuronal Ca2+ levels during all volitional whisking events 1½<3s long without (*dark blue*: All synaptic ROIs, *light blue*: synaptic ROIs within <15µm and 15-30µm from PA, *blue*: synaptic ROIs within 30-45µm and 45-60µm from PA) and with sandpaper (*orange*). Outline of 1 s.e.m. <u>Bottom</u>: Neuronal Ca2+ responses to volitional whisking with and without sandpaper averaged and statistically compared pr. FOV. Mixed model test, All: Max early p=0.088, Max late p=0.21, <15µm from PA: Max early p=0.021, Max late p=0.07, 15-30µm from PA: Max early p=0.022, Max late p=0.042, 30-45µm from PA: Max early p=0.79, Max late p=0.70, 45-60µm from PA: Max early p=0.13, Max late p=0.24, N= 19 FOVs in 7 mice. **F)** <u>Top</u>: Average astrocytic Ca2+ levels during all volitional whisking events 1½<3s long without sandpaper (*dark green*: All ROIs, *light green*: ROIs within <15µm and 15-30µm of PA, *green*: ROIs within 30-45µm and 45-60µm of PA) and with sandpaper (*orange*). Outline of 1 s.e.m. <u>Bottom</u>: Astrocytic Ca2+ responses to volitional whisking with and without sandpaper averaged and statistically compared pr. FOV. Mixed model test, All: Max early p=0.077, Max late: p=0.049, <15µm from PA: Max early: p=0.00471, Max late p=0.0154, 15-30µm from PA: Max early p=0.0073, Max late p=0.0069, 30-45µm from PA: Max early p=0.88, Max late p=0.51, 45-60µm from PA: Max early p=0.71, Max late p=0.63, N=20 FOVs in 16 mice **G)** Average 2-photon image from volitional whisking events without (above) and with (bottom) sandpaper. Ca2+ activity normalized to s.d. values per pixel over time (*pseudocolor*), before, at whisking onset, and in the first 5 seconds after. Overlayed image of vessels (*grey*), the red scale bar shows the dilation of PA (*dark red*) and 1^st^ order capillary (*light red*).

**Supplemental Figure S10:**
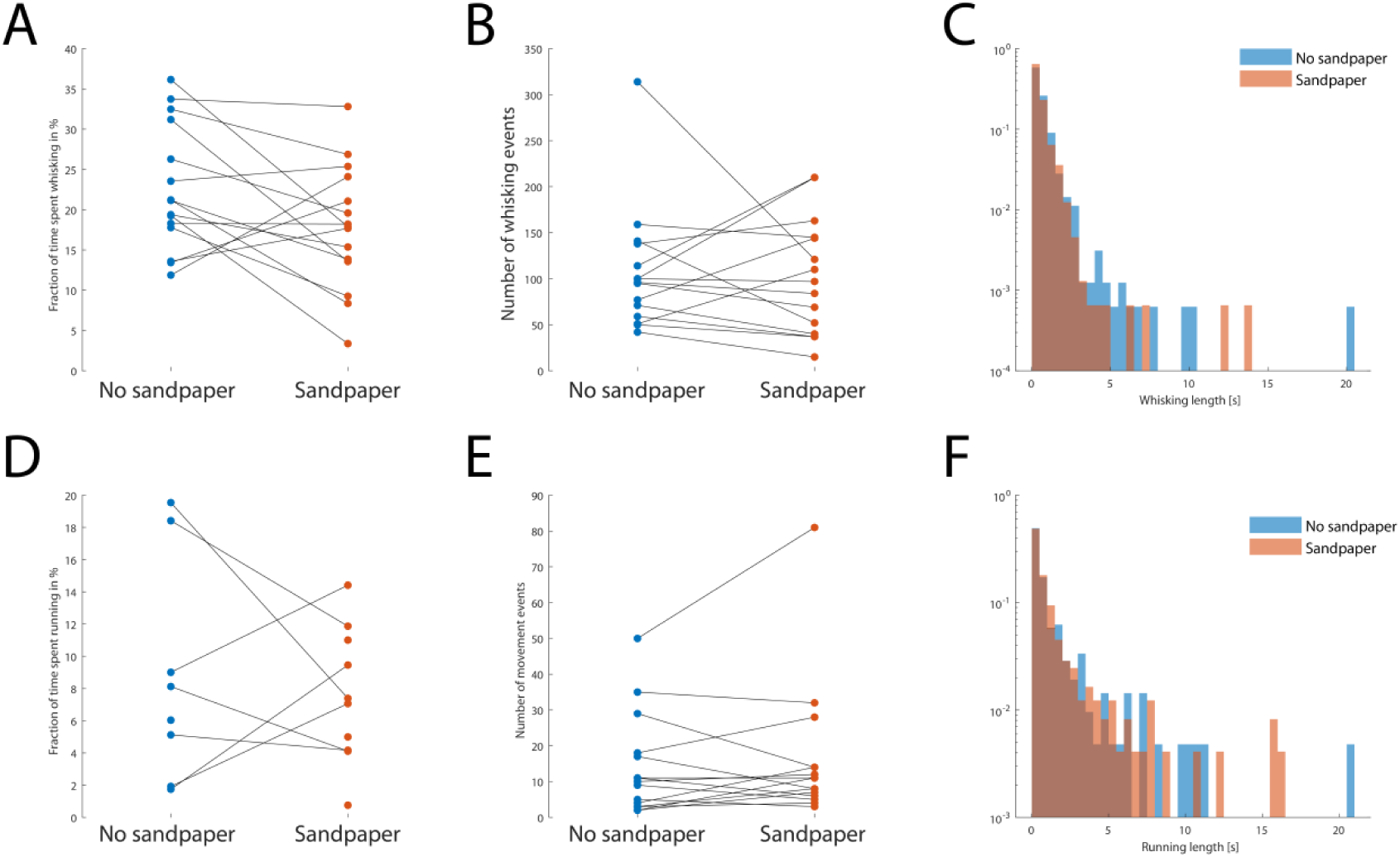
The presence of sandpaper did not influence animal behavior with regard to whisking or locomotion activity. **A)** Estimated per mouse, the presence of sandpaper did not change the fraction of time spent whisking. **B)** Nor did it influence the number of whisking events. **C)** Or the distribution of duration of whisking. **D)** Similarly, the sandpaper did not affect how much time the mouse spent running, **E)** how many locomotion events it had, **F)** or the distribution of the duration of the locomotions.

**Supplemental figure S11:**
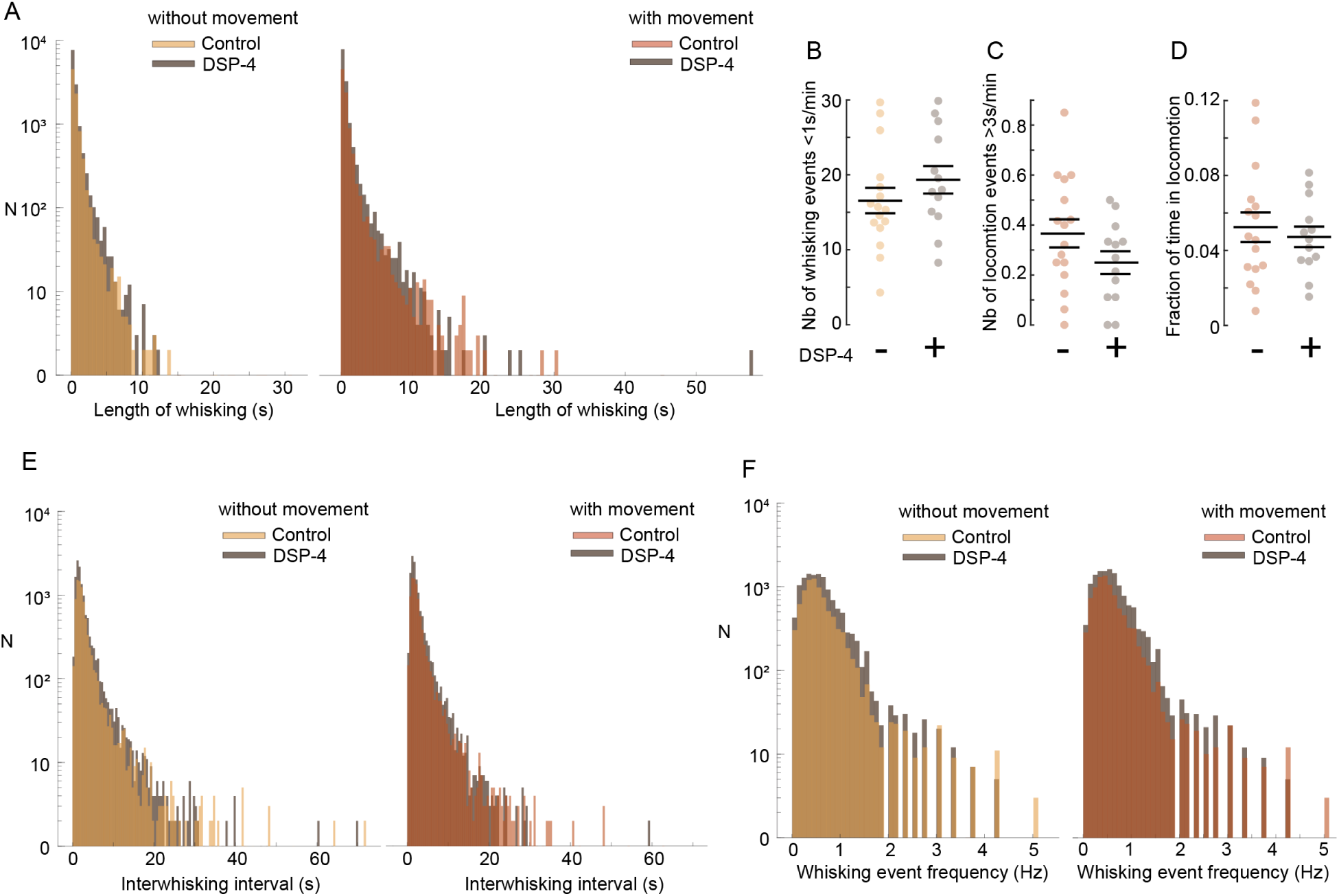
DSP-4 effect on whisking and locomotion behavior. **A)** Histograms showing the distributions of whisking events, including whisking during locomotion (*yellow*) and excluding locomotion-associated whisking (*red*). Whisking events longer than 5 seconds were very rare in the absence of running. **B)** We found no effect of DSP-4 on the fraction of time each animal spent moving. T-test p=0.61, N=16 mice. **C)** We found no effect of DSP-4 on how many locomotion events lasting longer than 3s could be detected in each animal. Mann-Whitney test, p=0.17, N= 16 mice. **D)** We found no effect of DSP-4 on how many short whisking events (<1s) could be detected in each animal. Mann-Whitney test, p=0.25, N= 16 mice. **E)** The time interval between two consecutive whisking events was usually quite short with the mode being 2 seconds. **F)** Taking the reciprocal of the interwhisk interval yields a whisking *frequency* with a mode of 0.5Hz.

**Supplemental figure S12:**
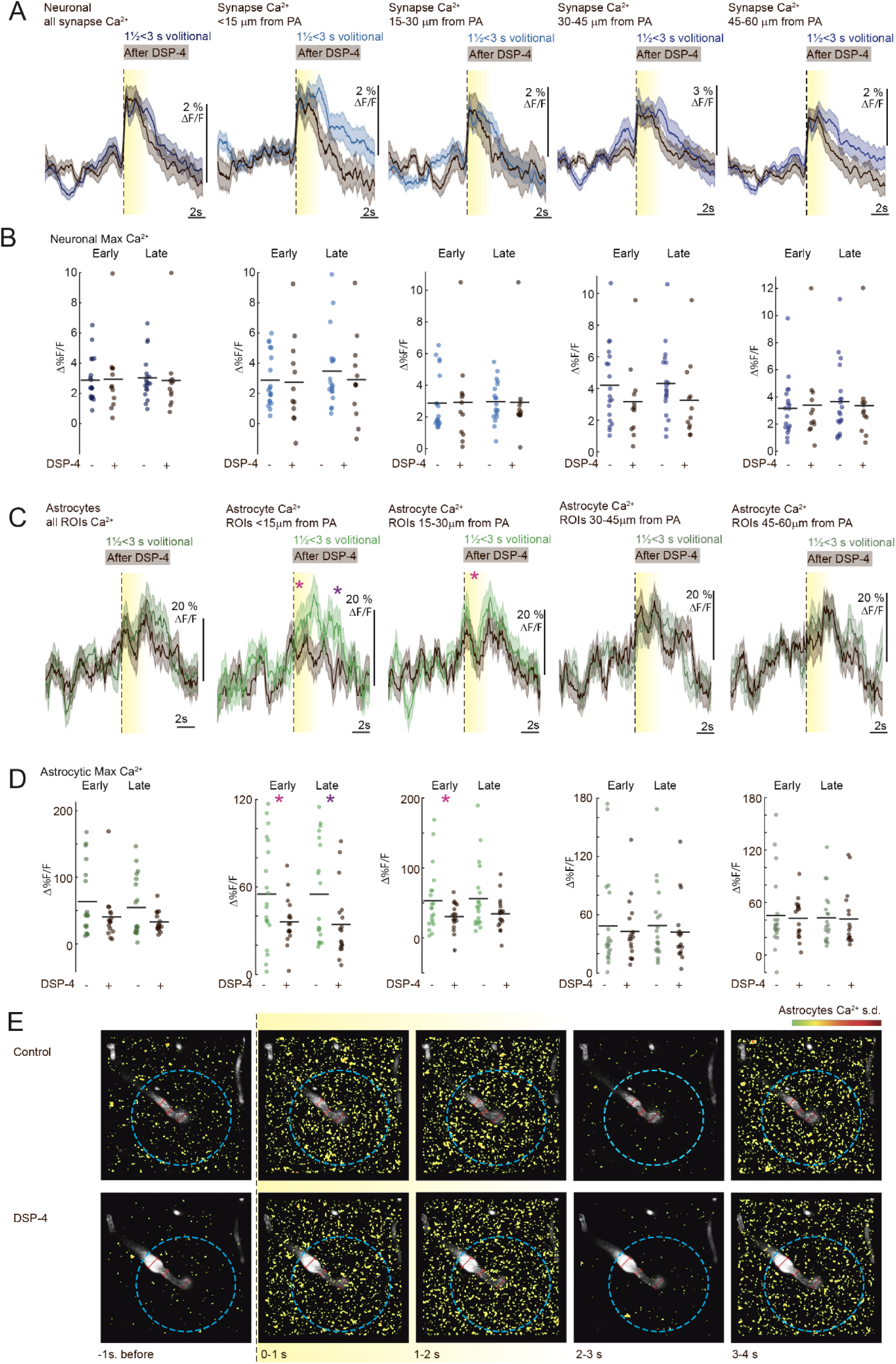
NA effect on volitional whisking response in neuronal and astrocytic Ca2+ depend on distance from PA. **A)** In no zones did the DSP-4 treatment affect the neuronal Ca2+ response to volitional whisking. Neuronal Ca2+ averaged from synaptic ROIs during volitional whisking events lasting between 1½-3 seconds. *Dark blue*: All ROIs, *Light blue*: Synaptic ROIs within <15µm and 15-30 µm of PA, *Blue*: Synaptic ROIs within 30-45µm and 45-60 µm of PA, *Black*: after DSP-4 treatment. Outline of 1 s.e.m. **B)** Quantification of the average effect of the DSP-4 treatment on neuronal Ca2+ in ROIs pr FOV. Mixed model, All ROIs: Early p=0.88, Late p=0.82, within 15µm of PA: early p=0.82, late p=0.57, 15-30 µm of PA: Early p=0.98. Late p=0.92, 30-45µm of PA: Early p=0.23 Late p=0.20 and 45-60 µm of PA: Early p=0.82. Late p=0.78, N=19 FOV in 7 mice. **C)** In the zones closest to the vessel, DSP-4 reduced astrocytic Ca2+ response to volitional whisking. Average astrocytic Ca2+ in ROIs during volitional whisking events lasting between 1½-3 seconds. *Dark green*: All ROIs, *light green*: ROIs within <15µm and 15-30µm of PA, *green*: ROIs within 30-45µm and 45-60µm of PA. Outline of 1 s.e.m. **D)** Quantification of the average effect of the DSP-4 treatment on astrocytic Ca2+ in ROIs pr FOV. Mixed model, All ROIs: Early Peak p=0.12, Late peak p=0.061, within 15µm of PA: Early peak p=0.0216, Late Peak: p=0.013, 15-30µm of PA: Early peak p=0.043, Late Peak: p=0.084, 30-45µm of PA: Early peak p=0.66, Late Peak: p=0.52, 45-60µm of PA: Early peak p=0.75, Late Peak: p=0.88, N=20 FOV in 16 mice. **E)** Average 2-photon image from volitional whisking events before (above) and following DSP-4 (below). Ca2+ activity normalized to s.d. values per pixel over time (*pseudocolor*), before, at whisking onset, and 3 seconds after. In the overlayed greyscale image of vessels, the red scale bar shows the dilation of PA (*dark red*) and 1^st^ order capillary (*light red*). *Blue dashed line* shows a 30µm distance from PA.

**Supplemental figure S13:**
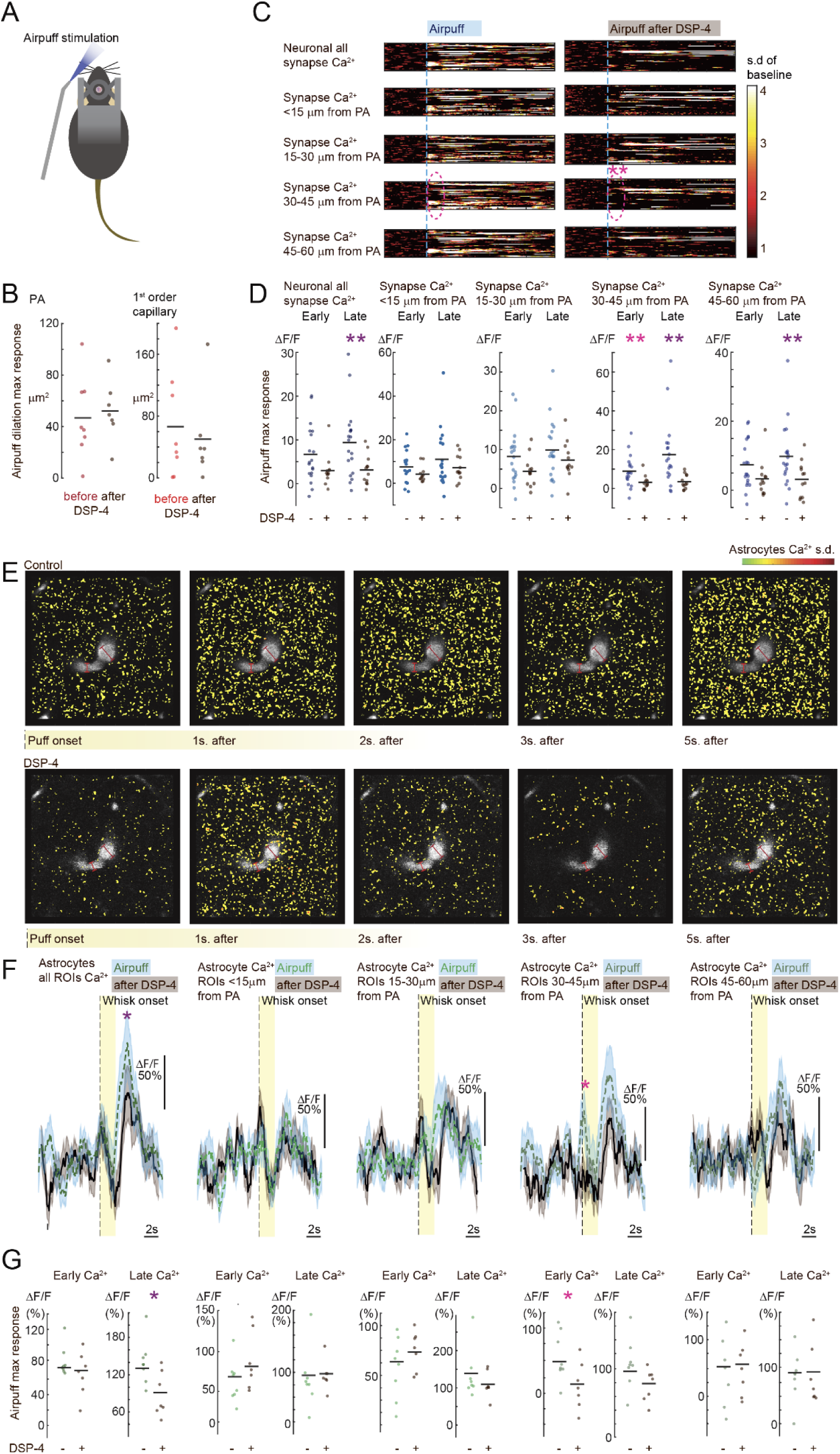
Neuronal and astrocytic responses to air-puff evoked whisking were reduced after DSP-4, but not within 30 µm of the vessel. **A)** Cartoon of the evoked whisker stimulation using air-puff. **B)** Quantification of the lack of effect of the DSP-4 treatment on the PA and 1^st^ order capillary dilations evoked by air-puff stimulation of the whiskers. Responses averaged pr FOV, mean shown as a *black line*. Paired t-test. PA: p=0.718, 1^st^ order capillary: p=0.632 N=8 FOV in 6 mice. **C)** Neuronal Ca2+ activity averaged from all synaptic ROIs dependent on distance from the center of the PA. Ca2+ levels normalized to s.d. values relative to 10s baseline shown in pseudocolor scale. Each air-puff evoked event aligned to whisking onset (*light blue line*). <u>Left:</u> before, <u>Right</u>: after DSP-4 treatment. *Pink circles, dashed line* mark the lack of early response in the synaptic ROIs further from the vessel. **D)** Quantification of peak neuronal Ca2+ response amplitude in activity averaged from synaptic ROIs dependent on distance from the center of the PA. DSP-4 only significantly reduced the responses in the synaptic ROIs more than 30 µm from the PA. Neuronal Ca2+ averaged from synaptic ROIs during 2s air-puff evoked whisking. Paired t-test, All ROIs: Early peak p=0.052, Late peak p= 0.0053, ROIs<15µm: Early peak p=0.10, Late peak p= 0.26, ROIs 15-30µm: Early peak p=0.061, Late peak p= 0.25, ROIs 30-45µm: Early peak p=0.0088, Late peak p= 0.0022, ROIs 45-60µm: Early peak p=0.073, Late peak p= 0.015, N=19 FOV in 7 mice. **E)** Average 2-photon image from air puff evoked events before (above) and following DSP-4 (below). Ca2+ activity normalized to s.d. values per pixel over time (*pseudocolor*), up to 5 seconds after air puff. Overlayed image of vessels (*grey*), the red scale bar shows the dilation of PA (*dark red*) and 1^st^ order capillary (*light red*). **F)** Astrocytic Ca2+ averaged from all ROIs and sorted dependent on distance from PA center during 2s air-puff evoked whisking. Before the drug, *Dark green*: All ROIs, *light green*: ROIs within <15µm and 15-30µm of PA, *green*: ROIs within 30-45µm and 45-60µm of PA. *Dashed line*, 1 s.e.m. outline in *light blue*. After DSP-4: *black, solid line*, s.e.m. outline in *dark grey*. **G)** Quantification of the average effect of the DSP-4 treatment on astrocytic Ca2+ in ROIs pr. FOV. Paired t-test, All ROIs: Early peak p=0.24, Late peak p=0.018, within 15µm of PA: Early peak p=0.17, Late Peak p=0.90, 15-30µm of PA: Early peak p=0.19, Late Peak p=0.37, 30-45µm of PA: Early peak p=0.031, Late Peak p=0.086, 45-60µm of PA: Early peak p=0.91, Late Peak p=0.97, N=8 FOV in 6 mice.

**Supplemental figure S14:**
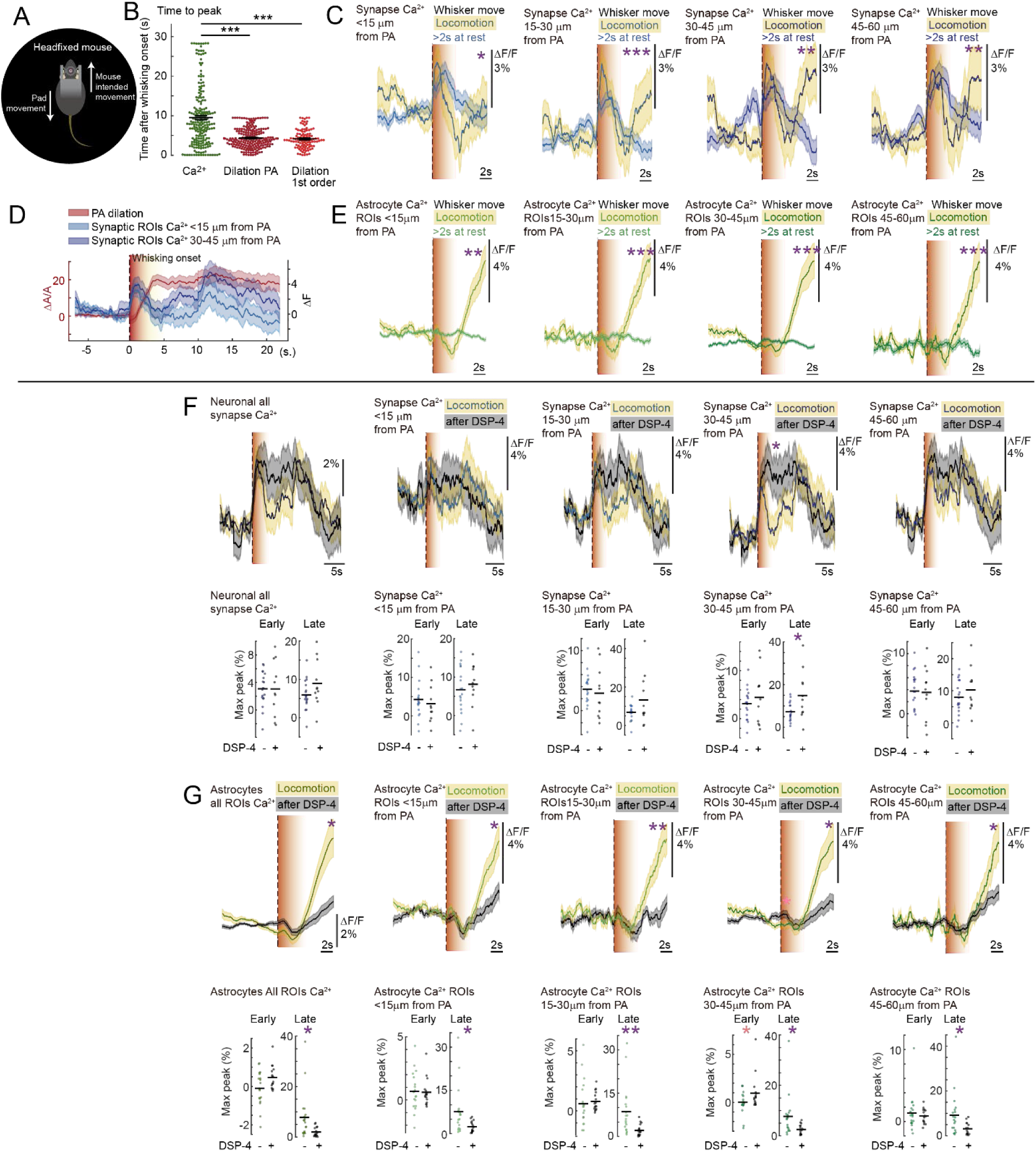
Neuronal and astrocytic Ca2+ responses to locomotion EXTRA. **A)** Cartoon of Neurotar system recording speed and direction of movement. **B)** Times to peak of activity from whisking onset associated with locomotion. Dilations peaked significantly faster than the astrocytic Ca^2+^ rises both for the PA (Mixed model, p = 3.33e-16) and the 1^st^ order capillary (Mixed model, p = 3.8e-11). **C)** Neuronal Ca2+ averaged from synaptic ROIs is dependent on distance from the center of the PA in >2s long whisking events during locomotion (s.e.m. outline *yellow*) compared to rest (s.e.m. outline *blue*). *Light blue*: Synaptic ROIs within <15µm and 15-30 µm of PA, *Blue*: Synaptic ROIs within 30-45µm and 45-60 µm of PA. A mixed model test of max amplitude showed increased response during locomotion but only late in the response. ROIs within 15µm of PA: Early: p=0.59 Late: p=0.021, 15-30 µm of PA: Early: p=0.87, Late: p=0.00064, 30-45µm of PA: Early: p=0.34 Late: p=0.0042 and 45-60 µm of PA: Early: p=0.63 Late: p=0.0093, N=18 FOV in 7 mice. **D)** Averages of all locomotion events aligned to the whisking onset. The *orange* area marks the period after whisking onset wherein locomotion began. PA vasodilation and Ca^2+^ activity in all neuronal synaptic ROIs <15µm (*light blue*) and from between 30-45µm distance to the PA (*dark blue*). N=18 FOV in 7 mice. **E)** Average astrocytic Ca2+ from all ROIs is dependent on distance from the center of the PA in >2s long whisking events during locomotion (s.e.m. outline *yellow*) compared to the rest (s.e.m. outline *green*). *Light green*: ROIs within <15µm and 15-30µm of PA, *green*: ROIs within 30-45µm and 45-60µm of PA. A mixed model test of max amplitude showed increased response during locomotion but only late in the response. ROIs within 15µm of PA: Early peak: p = 0.54, Late peak: p=0.0019, 15-30µm of PA: Early peak p=0.82, Late Peak: p=0.00021, 30-45µm of PA: Early peak p=0.18, Late Peak: p=0.00085, 45-60µm of PA: Early peak p=0.96, Late Peak: p=0.00079, N=19 FOV in 16 mice. **F)** <u>Top</u>: Neuronal Ca2+ averaged from all synaptic ROIS or dependent on distance from PA during locomotion. Control, *yellow* s.e.m. outline, *Dark blue*: All ROIs, *Light blue*: Synaptic ROIs within <15µm and 15-30 µm of PA, *Blue*: Synaptic ROIs within 30-45µm and 45-60 µm of PA. After DSP-4: Black line, *grey* s.e.m. outline. <u>Bottom</u>: Quantification of max amplitudes Ca2+ response to locomotion averaged pr FOV. Mixed model test. All ROIs: Early p = 0.68, Late p=0.12, within 15µm of PA: Early peak p=0.39, Late Peak p=0.55, 15-30µm of PA: Early peak p=0.48, Late Peak p=0.055, 30-45µm of PA: Early peak p=0.34, Late Peak p=0.012, 45-60µm of PA: Early peak p=0.74, Late Peak p=0.27, Control N=18 FOV, DSP-4 N=11 FOV in 7 mice. **G)** <u>Top</u>: Astrocytic Ca2+ averaged from all ROIS or dependent on distance from PA during locomotion. Control, *yellow* s.e.m. outline, *Dark green*: All ROIs, *Light green*: ROIs within <15µm and 15-30 µm of PA, *Green*: ROIs within 30-45µm and 45-60 µm of PA. After DSP-4: *Black line*, *grey* s.e.m. outline. <u>Bottom</u>: Quantification of max amplitudes Ca2+ response to locomotion averaged pr FOV. Mixed model test. All ROIs: Early p = 0.22, Late p=0.017, within 15µm of PA: Early peak p=0.86, Late Peak p=0.014, 15-30µm of PA: Early peak p=0.96, Late Peak p=0.0034, 30-45µm of PA: Early peak p=0.026, Late Peak p=0.015, 45-60µm of PA: Early peak p=0.67, Late Peak p=0.025, Control N=19 FOV in 16 mice, DSP-4 N=18 FOV in 13 mice.

**Supplemental figure S15:**
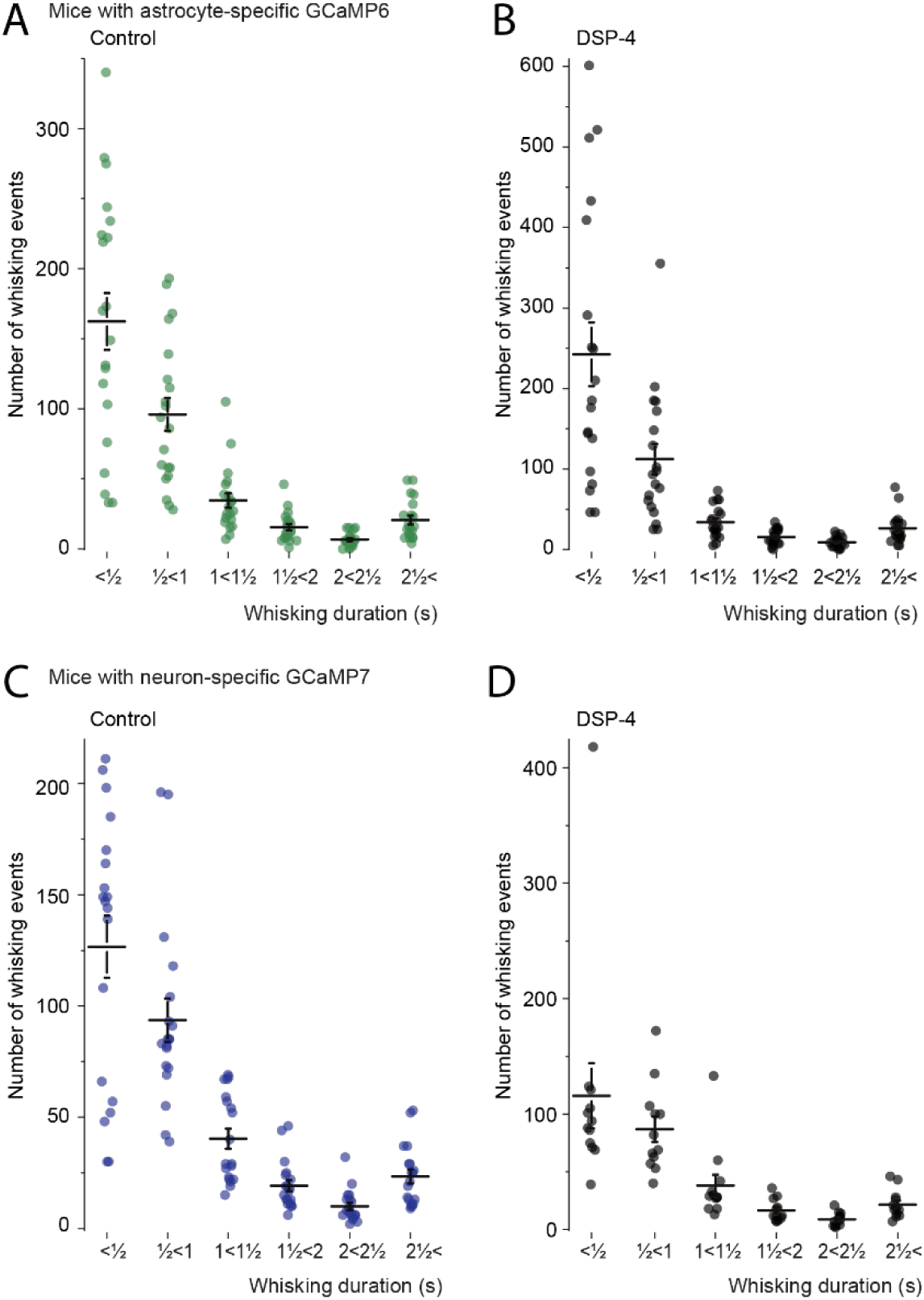
Number of volitional whisking events per FOV. **A)** Based on whisking duration in non-treated mice expressing the astrocyte specific Ca2+ indicator. N=20 FOV in 16 mice. **B)** Based on whisking duration in DSP-4 treated mice expressing the astrocyte specific Ca2+ indicator. N=19 FOV in 14 mice. **C)** Based on whisking duration in non-treated mice expressing the neuron specific Ca2+ indicator. N=19 FOV in 7 mice. **D)** Based on whisking duration in DSP-4 treated mice expressing the neuron specific Ca2+ indicator. N=19 FOV in 7 mice.

